# *Candida albicans* reprograms host inosine metabolism to drive immunosuppressive macrophage polarization and gastric cancer carcinogenesis

**DOI:** 10.64898/2026.05.27.728170

**Authors:** Jinshui Tan, Yubo Xiong, Ao Cheng, Chen-song Zhang, Mengya Zhong, Jingsong Ma, Jiabao Zhao, Yifan Zhuang, Wei Wang, Guangchao Pan, Zeyang Lin, Shengyi Zhou, Huiwen Zhou, Guoqiang Su, Xuehui Hong

## Abstract

Gastric cancer (GC) is strongly associated with changes in the gastric microbiome. However, the direct contribution of individual pathogenic species to tumor initiation and progression remains poorly defined. Here, we identified *Candida albicans* (*C. albicans*), a fungus that ranks as the most significantly enriched taxon in GC tissues, can directly promote gastric carcinogenesis. We found that when *C*. *albicans* alone was colonized into the gastric mucosa of mice, it induced the complete sequence of GC-associated preneoplastic lesions, including chronic gastritis, mucinous metaplasia, and atypical hyperplasia, with 16% of colonized animals progressing to low-grade intraepithelial dysplasia. In *K19-C2mE* transgenic mice, a model of spontaneous GC formation, infection with *C*. *albicans* significantly accelerated GC formation, with synergistic effects observed upon co-infection with the well-known gastric carcinogen, *Helicobacter pylori*. Mechanistically, *C*. *albicans* upregulated inosine levels in both gastric tissues and circulation. This inosine promoted polarization of macrophages toward an M2-like phenotype by activating the adenosine A2a receptor (A2aR), creats an immunosuppressive microenvironment that protects GC cells from the tumoricidal effects of anti-tumor immunity. Depletion of M2 macrophages or pharmacological inhibition of A2aR attenuated *C*. *albicans*-mediated tumorigenesis. Clinically, elevated inosine levels correlated with prior *C*. *albicans* exposure and poorer outcomes of GC patients. Our findings establish *C*. *albicans* as a direct pro-tumorigenic pathogen in GC and highlight the inosine-A2aR-M2 macrophage axis as a promising target for therapeutic intervention.

## Introduction

GC is the fourth leading cause of cancer-related deaths worldwide, with a dismal five-year survival rate below 30%, largely attributable to late diagnosis and limited treatment options^1,2^. It has been well established that the gastric microbiota is a key player in carcinogenesis, with *Helicobacter pylori* (*H. pylori*) recognized as a class I carcinogen and a major risk factor^3^. In addition to *H. pylori*, other microbes, including *Streptococcus anginosus* (*S. anginosus*), *Fusobacterium*, *Propionibacterium acnes*, *Prevotella intermedia*, *Neisseria* and virus species like Epstein-Barr virus (EBV) have been associated with poorer outcomes in GC^4–10^. However, despite these advances, the etiological contribution of individual microbes to a significant proportion of GC cases remains unclear. For instance, although *H*. *pylori* infection is widespread, it is detected in only about 35% of GC patients, and epidemiologic studies suggest that only 3% of infected individuals progress to preneoplastic lesions^11,12^. Similarly, *S*. *anginosus* has been identified as an independent driver in at most 16% of GC cases^4,13^, while EBV is implicated in approximately 10% of cases^14^. This indicates that gastric carcinogenesis is likely driven by a complex interplay of microbial and host factors, and that dominant, broadly prevalent drivers still remain to be fully identified.

In addition to bacteria and viruses, fungi represent an integral but understudied component of the gastric microbiome. In our previous study, we analyzed the fungal communities in samples from both GC patients and non-cancerous controls. We found that *C*. *albicans* is the most significantly enriched taxon of fungi in GC tissues, accounting for approximately 22% of the total fungal abundance^15^. This finding was further supported by a follow-up meta-analysis using data from the TCGA database, which demonstrated a higher proportion of *C*. *albicans* in GC patients, and exhibits a strong positive correlation with tumor stage among^16^. Consequently, *C*. *albicans* has been identified as the first fungal pathogen linked to GC. In particular, the high prevalence and stage-associated enrichment suggest that *C*. *albicans* may play a broader and more direct role in gastric carcinogenesis than previously recognized.

Here, we demonstrate that *C. albicans* functions as a direct driver of GC. Using specific pathogen-free (SPF) and germ-free mice, we show that monocolonization with *C. albicans* is sufficient to induce GC-related preneoplastic progression and, in 16% cases, low-grade intraepithelial dysplasia. In a transgenic model of spontaneous GC (*K19-C2mE* mice), *C*. *albicans* accelerates tumor development to an extent comparable to that of *H*. *pylori* and can have enhanced effects when co-infected. Mechanistically, we identify a pathway through which *C. albicans* elevates inosine levels, thereby polarizing macrophages toward a tumor-promoting M2-like phenotype via the adenosine receptor A2aR. Clinically, the levels of inosine correlate with exposure to *C*. *albicans* and predict worse outcomes. Our work establishes *C*. *albicans* as a pro-tumorigenic gastric pathogen and reveals the inosine-A2aR-macrophage axis as a potential therapeutic target in GC.

## Results

### *C. albicans* triggers gastric tumorigenesis in mice

We gavaged wildtype C57BL/6 SPF mice with *C. albicans* (1 x 10^8^ CFU per dose) every three days (following a dose commonly used in mouse studies^17,18^), allowing sustained colonization of this fungus in the gastric mucosa (**Figures S1A-S1C**) and evaluated the phenotype at 3, 6, 9, and 12 months post-infection (**Figure 1A**). The successful colonization of *C. albicans* in the gastric mucosa was validated using FISH in situ (**Figure 1B**), as well as the bacterial culture of freshly harvested stomach tissues followed by MALDI-TOF MS and PCR analysis (**Figures S1D-S1G; Table S1**). Compared to the PDB□broth□gavaged controls, *C*. *albicans*□infected mice developed chronic gastric inflammation by 3 months (characterized by sustained lymphocyte infiltration and the appearance of submucosal lymphoid aggregates, and assessed by inflammation scores; **Figures 1C and 1D**), which progressed over the next 6 months and persisted through the 12-month mark (**Figures 1C and 1D**). Parietal cell atrophy was evident after 9 months of infection and continued to develop from 9 to 12 months post-infection (indicated by atrophy scores; **Figures 2A and 2B**). Consistent with the induction of gastric atrophy, *C. albicans* infection increased gastric pH after 12 months compared with the PDB control group (**Figure S1H**). In addition, we observed a 1.5-fold increase in cell proliferation rates (assessed by Ki-67-positive staining) in the gastric mucosa from mice after 6 months of infection (**Figures 2C and 2D**), which became more pronounced, reaching approximately 2-fold after 12 months of infection (**Figures 2C and 2D**). Consistently, we also observed a higher incidence of mucinous metaplasia in the mice after 12 months of *C. albicans* infection (evidenced by Alcian blue staining; **Figures 2E and 2F**). Moreover, about 16% of the mice exhibited low-grade intraepithelial dysplasia (**Figures 2G and 2H**). These data suggested that the colonization of *C. albicans* is sufficient to initiate a complete histopathological sequence of GC-associated preneoplastic lesions.

**Figure 1.**
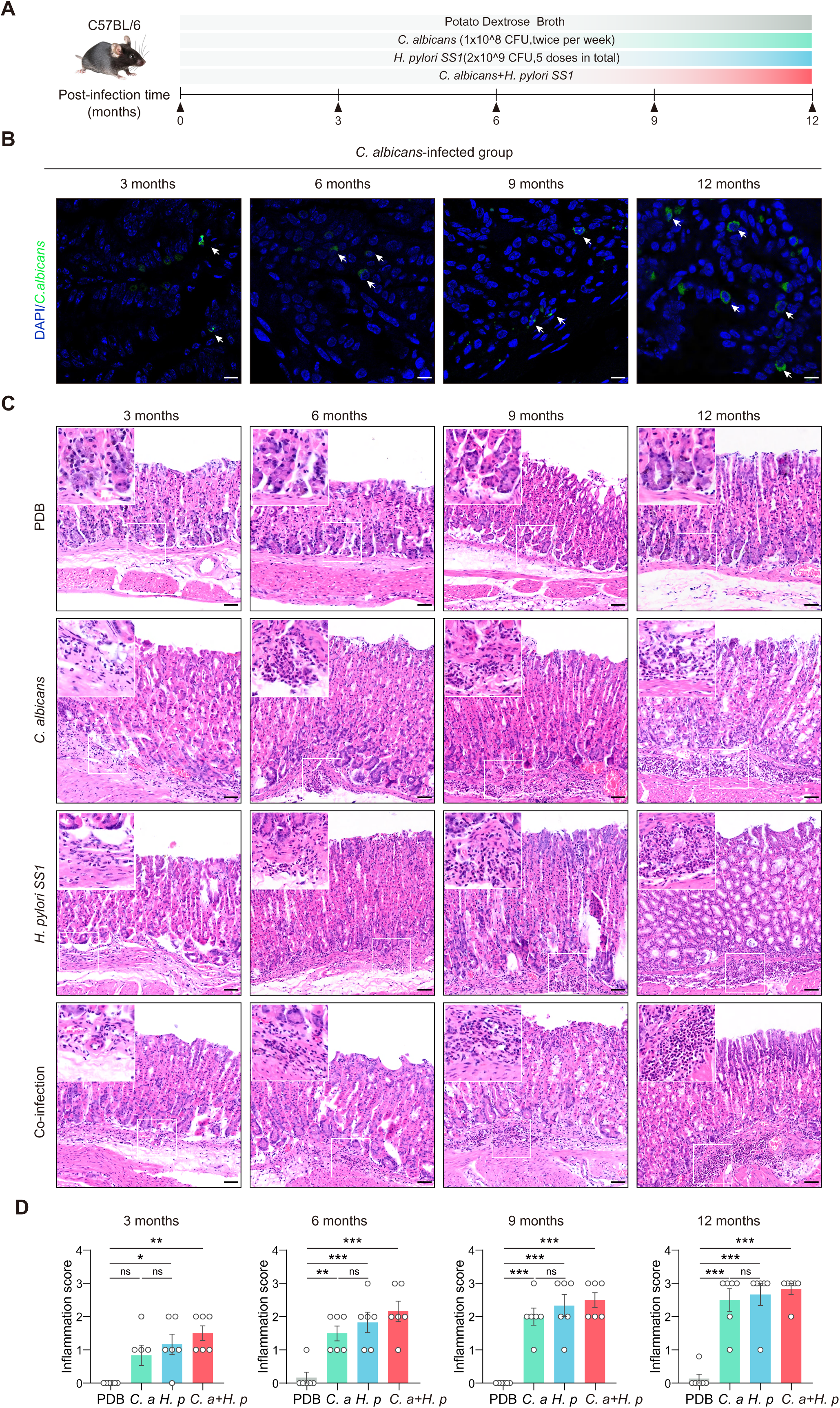
*Candida albicans* induces chronic gastritis in mice. (A) C57BL/6 male mice were orally administered PDB, *C. albicans* (*C. a*), *H. pylori* (*H. p*), or co-administration (*C. a* + *H. p*) for 3, 6, 9, and 12 consecutive months (n = 6/group/time point) (B) Representative FISH-stained stomach sections from mice infected with *C. albicans* at 3, 6, 9, and 12 months post-infection; scale bars, 10 μm. (C) Representative H&E-stained images (inflammation) of stomach sections (3, 6, 9, and 12 months post-infection) from PDB, *C. albicans* (*C. a*), *H. pylori* (*H. p*), or co -administration (*C. a* + *H. p*); scale bars, 100μm. (D) Histological inflammation scores of gastric sections (3, 6, 9, and 12 months post-infection) in the indicated groups. Data were presented as mean ± SEM. Each point represents one subject. Kruskal-Wallis test (D) was performed to assess differences among groups, ns, nonsignificant, * p < 0.05, ** p < 0.01, *** p < 0.001.

**Figure 2.**
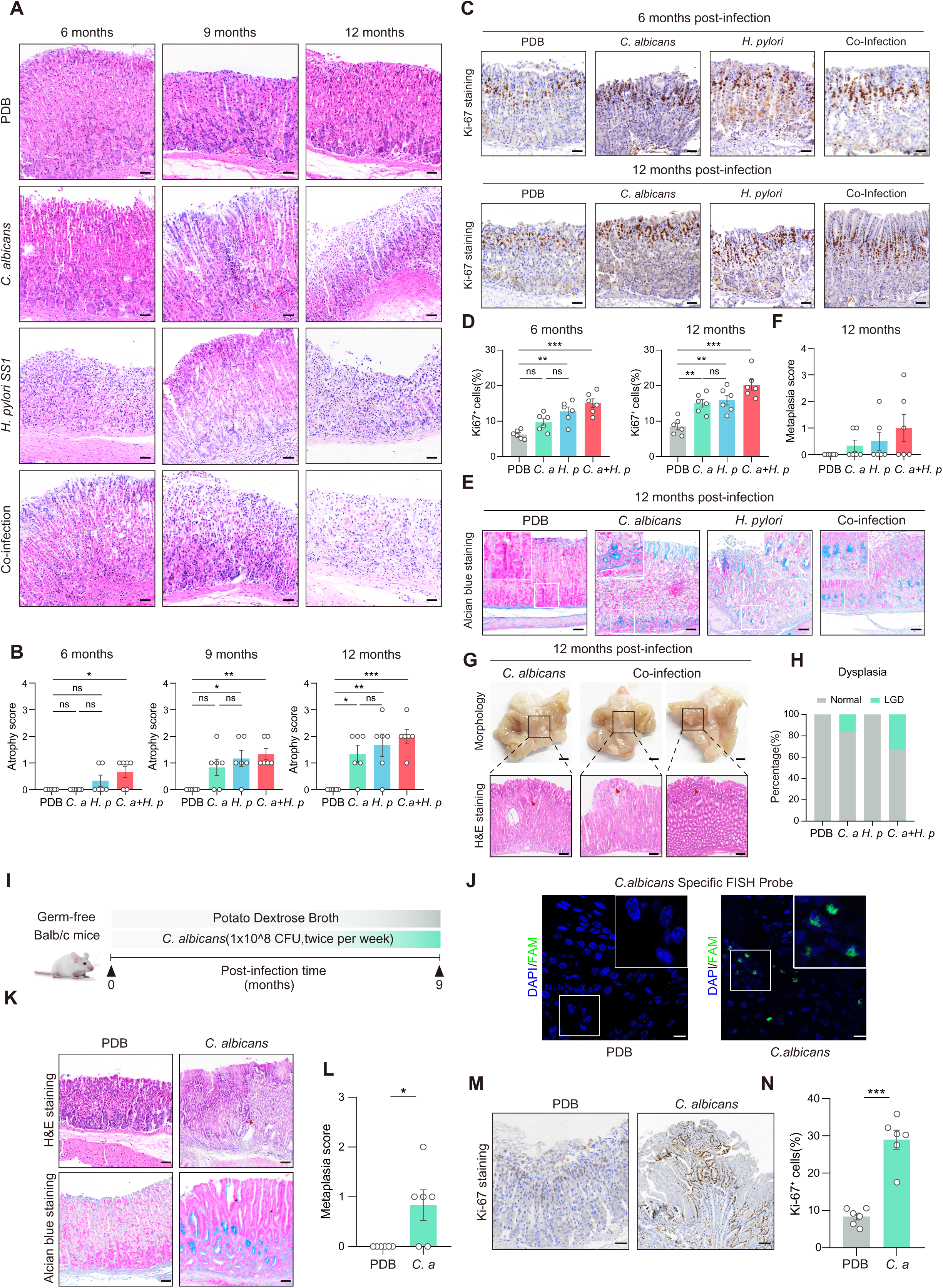
*Candida albicans* promotes preneoplastic lesions in mice. (A) Representative H&E-stained images (atrophy) of stomach sections (6, 9, and 12 months post-infection) from PDB, *C. albicans* (*C. a*), *H. pylori* (*H. p*), or co -administration (*C. a* + *H. p*); scale bars, 100μm. (B) Histological atrophy scores of gastric sections (6, 9, and 12 months post-infection) in the indicated groups. (C and D) Representative images of Ki-67 staining and proliferation rates in the indicated groups at 6 (C) and 12months (D) post-infection. scale bars, 100μm. (E and F) Representative alcian blue-stained images (mucinous metaplasia, E) and metaplasia score (F) of stomach sections at 12 months from PDB, *C. albicans* (*C. a*), *H. pylori* (*H. p*), or co -administration (*C. a* + *H. p*); scale bars, 100μm. (G) Representative morphology image and H&E-stained images(dysplasia) of stomach sections at 12 months from *C. albicans* (*C. a*) and co -administration (*C. a* + *H. p*); scale bars, 50mm (morphology), 100μm (H&E). (H) The proportion of dysplasia in the indicated groups at 12 months. (I) Germ-free male mice were orally administered PDB or *C. albicans* (n = 6/group) twice weekly for 9 consecutive months. (J) Representative FISH-stained image of stomach sections infected with *C. albicans* at 9 months post-infection; scale bars, 10 μm. (K) Representative H&E-stained images and alcian blue-stained images of stomach sections infected with PBD or *C. albicans* at 9 months post-infection; scale bars, 100 μm. (L) Metaplasia score of stomach sections infected with PBD or *C. albicans* at 9 months post-infection. (M and N) Representative images of Ki-67 staining and proliferation rates in the indicated groups at 9 months post-infection. scale bars, 100μm. Data were presented as mean ± SEM. Each point represents one subject. Kruskal-Wallis test (B and F), One-way ANOVA (D) and Student’s t test (L and N) were used to assess the statistical significance between groups, ns, nonsignificant, * p < 0.05, ** p < 0.01, *** p < 0.001.

To establish the causal role of *C. albicans* in gastric tumorigenesis without the influence of a complex microbiome, we introduced *C. albicans* into germ-free mice (**Figure 2I**) (see validation data for the colonization of *C. albicans* in **Figure 2J, Figures S1I and S1J; Table S1**). After 9 months of infection, we observed a significant increase in mucinous metaplasia (**Figures 2K and 2L**), along with gastric alkalization (**Figure S1K**), heightened cell proliferation (**Figures 2M and 2N**), confirming that *C*. *albicans* itself is sufficient to drive preneoplastic changes of GC.

The pronounced effects of *C. albicans* on gastric tumorigenesis are reminiscent of those caused by the well-established gastric carcinogen, *H*. *pylori*^3^, and prompted us to compare them. As shown in **Figures 1C and 1D**, **Figures 2A-2D**, *C. albicans* infection led to gastric inflammation, parietal cell atrophy, and increased proliferation of gastric mucosa cells, similar to the effects observed with *H*. *pylori* at each time point examined. While *H*. *pylori* infection resulted in more severe mucinous metaplasia than that of *C. albicans*, it did not lead to any intraepithelial dysplasia in the infected mice (**Figures 2E–2H**; also consistent with previous studies^4^. We also co-infected the mice with a mixture of the two pathogens, and observed synergistic effects across all examined pathological outcomes (**Figures 1C and 1D**, **Figures 2A–2F**). Notably, over 30% of the infected mice exhibited low-grade intraepithelial dysplasia (**Figures 2G and 2H**). Therefore, *C. albicans* showed comparable effectiveness in gastric tumorigenesis to *H*. *pylori* and acted synergistically with it to promote gastric lesions.

We next determined the roles of *C. albicans* in the formation of GC tumors. For this purpose, we co-cultured PDB, *Saccharomyces cerevisiae* (*S. cerevisiae*), and *C. albicans* with gastric cancer cells, and found that *C. albicans* did not stimulate tumor cell proliferation *in vitro* (**Figures S2A and S2B**). Subsequently, we utilized the *K19*-*C2mE* transgenic mice, which exhibit accelerated gastric inflammation and early development of dysplasia, and later formation of gastric tumors^19^, thereby allowing the spontaneous development of GC tumors within their lifespans when exposed to carcinogens such as *H*. *pylori*. We first determined the optimal dosing regimen for *C. albicans* in *K19-C2mE* mice. Through dose titration and qPCR assessment of gastric colonization, we determined that a dose of 1 × 10□ CFU, which we used in wildtype C57BL/6 mice, resulted in a level of *C*. *albicans* colonization comparable to that found in patient tissues (**Figure S2C**). Consequently, we selected this dosage for all subsequent experiments. We found that after 2 months of *C*. *albicans* infection (**Figure 3A**) (see validation data in **Figures S2D and S2E; Table S1**), GC tumors developed, which was characterized by both high-grade dysplasia (HGD) and well-differentiated adenocarcinoma (WDA), as evidenced by H&E staining (**Figures 3C and 3D**), and accompanied by gastric alkalization (**Figure S2F**). In contrast, the control group, which received *S. cerevisiae* or PDB medium gavage, only the low-grade dysplasia (LGD) was displayed, with no signs of HGD or WDA (**Figures 3C and 3D**). We also found that *C. albicans* infection led to larger tumor volumes and higher tumor numbers (**Figure 3E**), which correlated with increased proliferation of gastric mucosa cells (**Figure 3F**). This effect on GC development was comparable to that of *H*. *pylori*, which served as a positive control in the *K19*-*C2mE* mice (**Figures 3A-3F**).

**Figure 3.**
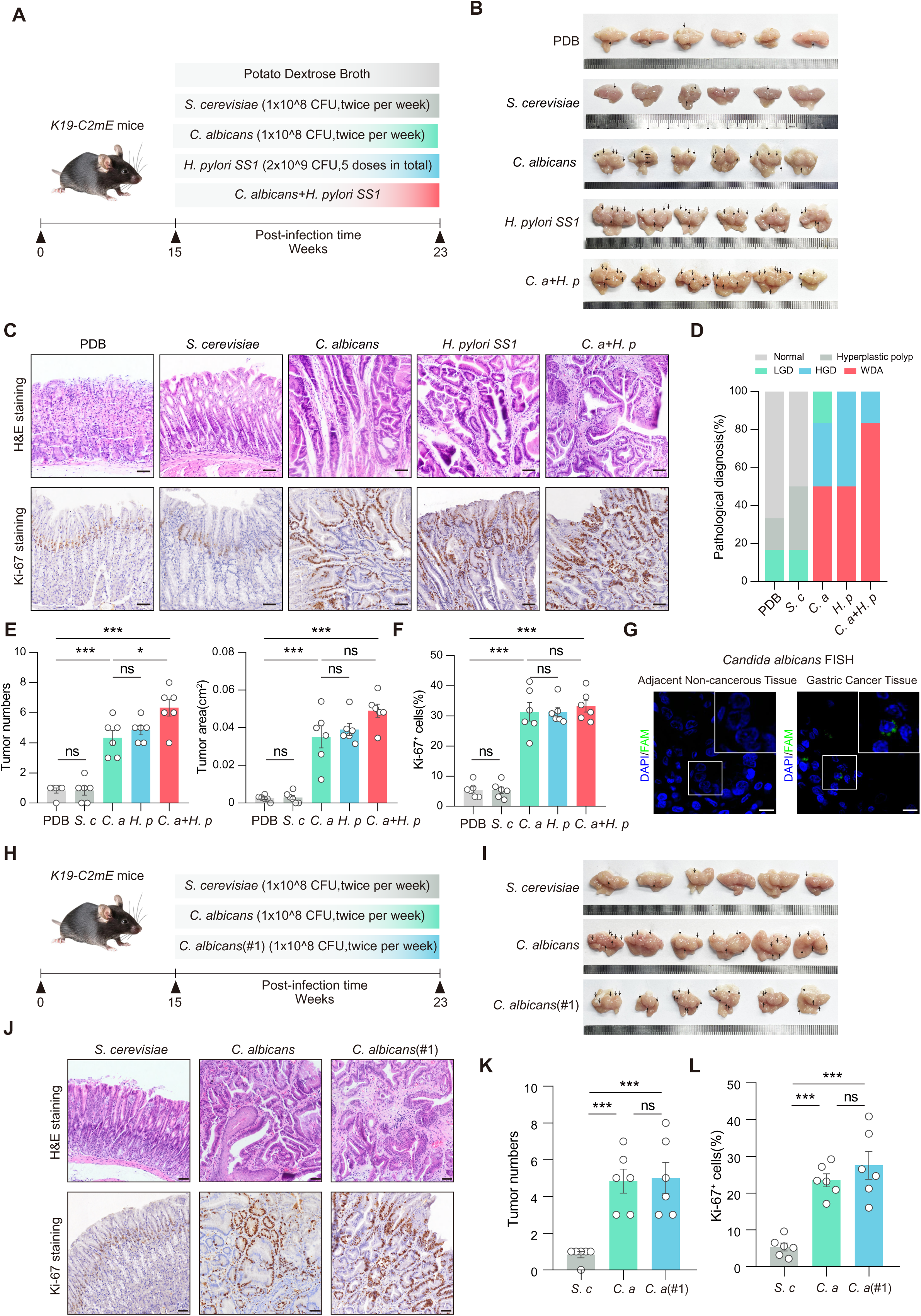
*Candida albicans* promotes the development of gastric cancer in mice. (A) *K19-C2mE* mice were orally administered PDB, *S. cerevisiae* (*S. c*), *C. albicans* (*C. a*), *H. pylori* (*H. p*), or co -administration (*C. a* + *H. p*) for 2 consecutive months (n = 6/group) (B) Representative morphology images of stomachs in the indicated groups; scale bars, 5 mm. (C) Representative H&E-stained and Ki-67-stained images of stomach sections from indicated groups; scale bars, 100μm. (D) The proportion of pathological diagnosis in the indicated groups. (E-F) Quantification of tumor numbers, tumor area (E) and Ki-67 proliferation rates(F) in the indicated groups. (G) Representative FISH-stained image of stomach sections in adjacent non-tumorous tissues and tumor samples from gastric cancer patients; scale bars, 10 μm. (H) *K19-C2mE* mice were orally administered *S. cerevisiae* (*S. c*), *C. albicans* (*C. a*), or clinical gastric cancer tissue isolated *C. albicans* (*C. a* #1) for 2 consecutive months (n = 6/group) (I) Representative morphology images of stomachs in the indicated groups; scale bars, 5 mm. (J) Representative H&E-stained and Ki-67-stained images of stomach sections from indicated groups; scale bars, 100μm. (K-L) Quantification of tumor numbers (K) and Ki-67 proliferation rates(L) in the indicated groups. Data were presented as mean ± SEM. Each point represents one subject. Kruskal-Wallis test (E/tumor numbers and K/tumor numbers) and One-way ANOVA (E/tumor area, F, K/tumor area, and L) were used to assess the statistical significance between groups, ns, nonsignificant, * p < 0.05, *** p < 0.001.

To further mimic the pathological characteristics of GC observed in patients, we isolated a *C. albicans* strain (*C. albicans*#1) from GC tissue obtained from a patient (validated in **Figure 3G, Table S1**). When this strain was introduced into the *K19*-*C2mE* mice, we observed a pro-tumor effect similar to that of the commercially available *C. albicans* strain (**Figures 3H–3L**). Treatment with the antifungal agent Amphotericin B (AmB) to eliminate *C. albicans* in the infected *K19*-*C2mE* mice impaired the pro-tumor effect associated with this fungus (**Figures S2G–S2L**). As a control, treatment with antibiotic cocktails that eliminate bacteria but not *C. albicans* showed no effect (**Figures S2G–S2L**). Although the germ-free *K19*-*C2mE* mice were unable to develop due to an unexpected technical issue, we can exclude dysbiosis of other bacterial flora caused by *C*. *albicans* infection as a contributing factor to the development of GC. This is supported by findings that *C*. *albicans* infection did not alter the composition of the gastric microbiome during the development of GC, as evidenced by microbial taxonomic profiling using ITS and 16S rDNA gene sequencing (**Figures S3A and S3B**), Shannon and Simpson indices (**Figures S3C and S3D**), principal coordinate analysis (PCoA; **Figures S3E and S3F**), and Linear discriminant analysis of effect size (LEfSe; **Figures S3G and S3H**). Results above indicate that *C. albicans* significantly promotes the formation of GC. Furthermore, we investigated the potential synergistic effect of *C. albicans* and *H*. *pylori* in promoting GC formation, and indeed observed a significantly higher number of tumors in co-infected mice (**Figures 3A–3F**).

### *C. albicans* elevates inosine levels for gastric tumorigenesis

We next explored the mechanisms through which *C. albicans* promotes gastric tumorigenesis. We found that gavage of heat-killed (HK; exposed to 95 °C for 2 h; see validation data in **Figure S3I**) *C. albicans* into the *K19*-*C2mE* mice did not induce GC (**Figures S2G–S2L**), indicating that merely having *C*. *albicans* present as an antigen is insufficient to trigger gastric tumorigenesis. In comparison, when culture supernatant from *C*. *albicans* co□cultured with gastric epithelial GES□1 cells was gavaged to *K19*-*C2mE* mice, tumor progression was significantly accelerated (**Figures S4A–S4E**). These results suggested that *C*. *albicans* must interact with host epithelial cells to generate tumor□promoting factor(s), likely metabolites.

To identify the responsible metabolite(s), we performed RNA□seq on the gastric tissues collected after being infected with *C*. *albicans* for 2 months, prior to the formation of GC (**Figure 4A, Figure S4F**). Pathway analysis revealed a marked enrichment for purine metabolism (**Figures 4B and 4C**). In addition, through untargeted metabolic mass spectrometry of gastric tissue from *C. albicans*-infected *K19*-*C2mE* mice collected at 2 months after infection (prior to GC formation) (**Figure 4D**), we identified 37 upregulated and 28 downregulated metabolites (**Figure 4E, Figure S4G; Table S2**). In serum from these mice, we also discovered 58 upregulated and 47 downregulated metabolites (**Figure 4F, Figure S4H; Table S2**). Among these metabolites, inosine was the unique shared metabolite found in altered metabolites in gastric tissue and serum, as determined by Venn diagram analysis (**Figure 4G**). The upregulation of inosine levels in gastric tissue and serum following *C*. *albicans* infection was further validated through the inosine assay kit (**Figures 4H and 4I, Figure S4I**). Importantly, *C. albicans* does not produce inosine *in vitro* within the culture system (**Figure 4J and Figure S4J**). However, its ability to stimulate inosine production was observed when co-cultured with GES-1 cells (**Figure 4J**). Notably, when *C*. *albicans* was removed from the co-culture, inosine secretion from GES-1 cells significantly decreased and was ultimately nearly eliminated (**Figure 4J**). Furthermore, the expression levels of the enzyme adenosine deaminase (ADA), responsible for converting adenosine to inosine^20^, were elevated in *C*. *albicans*-infected gastric cancer tissues and in *C*. *albicans*-infected GES-1 cells (**Figures 4K and 4L, Figure S4K**). Notably, we found that inosine elevation occurred exclusively in normal gastric epithelial cells post-*C*. *albicans* infection, as no elevation was observed in GC tumor cell lines such as AGS and MKN45 **(Figures S4L and S4M),** possibly due to the already low ADA levels in these cells (AGS and MKN45) (**Figure 4L**).

**Figure 4.**
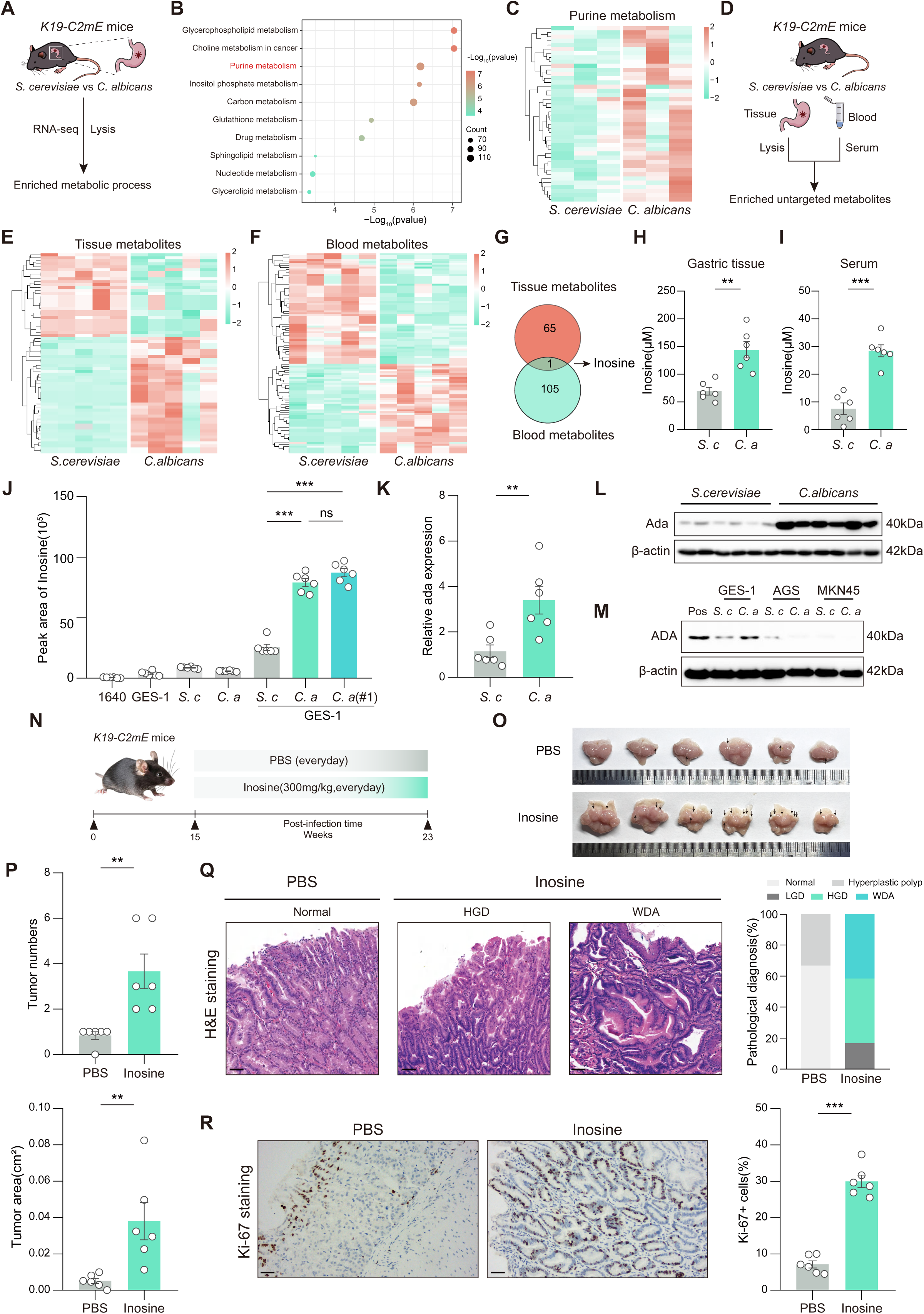
*Candida albicans* enhances inosine-induced purine nucleotide metabolism to promote gastric tumorigenesis. (A) Schematic diagram of the procedure for identifying biological processes of gastric tissue in *S. cerevisiae* or *C. albicans* infected mice (n = 3/group). (B) Differentially expressed genes were analyzed using KEGG enrichment for biological processes. (C) Heatmap of dysregulated genes in purine metabolism identified by RNA sequencing. (D) Schematic diagram of the procedure for metabolites of gastric tissue and serum in *S. cerevisiae* or *C. albicans* infected mice. (E-F) Heatmap of Significantly different metabolites (p < 0.05) of gastric tissue (E) and serum (F) identified by untargeted metabolomics (n = 6/group). (G) Venn diagram of shared metabolites with consistent changes in gastric tissue and serum. Inosine is the singular metabolite showing consistent patterns in both gastric tissue and serum. (H-I) Elisa assay of inosine concentration in gastric tissue (H) and serum (I). (J) The relative concentration of inosine in different treatment groups in GES-1 cells. (K) The mRNA expression of ADA in gastric tissue of *S. cerevisiae* or *C. albicans* infected *K19-C2mE* mice. (L) The protein expression of ADA in gastric tissue of *S. cerevisiae* or *C. albicans* infected *K19-C2mE* mice. (M) The protein expression of ADA in GES-1, AGS, MKN45 infected with *S. cerevisiae* or *C. albicans*. (N) *K19-C2mE* mice were orally administered PBS or inosine (300 mg/kg/day) for 8 weeks consecutively (n = 6/group). (O) Representative morphology images of stomachs in the indicated groups; scale bars, 5 mm. (P) Quantification of tumor numbers and tumor area in the indicated groups. (Q) Representative H&E-stained and proportion of pathological diagnosis of stomach sections from indicated groups; scale bars, 100μm. (R) Representative Ki-67-stained images and proliferation rate of stomach sections from indicated groups; scale bars, 100μm. Data were presented as mean ± SEM. Each point represents one subject. Mann-Whitney U test (P/tumor numbers), Student’s t-test (H, I, K, P/tumor area, and R) and one-way ANOVA test (J) were used to assess the statistical significance between groups, ** p < 0.01, *** p < 0.001.

We then asked whether inosine itself is sufficient to drive tumorigenesis. Daily gavage of *K19*□*C2mE* mice with inosine (300 mg/kg^21^, which led to the accumulation of inosine in the gastric epithelium similar to that observed after *C. albicans* infection, **Figures S4N and S4O**), or PBS as a control, for 8 consecutive weeks (**Figure 4M**) significantly elevates the number and volume of GC tumors, particularly well-differentiated adenocarcinomas (**Figures 4N–4Q**). Additionally, inosine increased the proliferation rates of gastric epithelial cells in these mice (**Figure 4R**). Therefore, *C*. *albicans* stimulated epithelial cells to secrete inosine, thereby contributing to gastric tumorigenesis.

### *C. albicans* promotes M2 macrophage polarization during gastric tumorigenesis

We therefore examined whether inosine may promote the proliferation of GC cells, and observed that, surprisingly, incubating inosine with AGS and MKN45 cells at varying concentrations and durations did not affect cell proliferation (**Figures S4P**). Inosine also did not affect the colony formation of these cells (**Figures S4Q and S4R**). Given that inosine and *C*. *albicans* can promote gastric tumorigenesis *in vivo*, we speculated that their tumor-promoting effects likely depend on the tumor microenvironment.

We thus reanalyzed our RNA-seq data from gastric tumors in *K19*-*C2mE* mice, and discovered that the *C*. *albicans*-infected group exhibited enrichment for macrophage-related pathways, including macrophage migration, differentiation, and activation (**Figure S5A**). Seq-ImmuCC analysis also indicated a significant increase in macrophages within the *C*. *albicans*-infected group (**Figure S5B**). To systemically determine whether *C*. *albicans* mediates a protumor effect via immune cells, we performed unbiased 10x single-cell RNA sequencing (scRNA-seq) analysis on gastric tumors from *K19*-*C2mE* mice infected with *C*. *albicans* (**Figure 5A**). As shown in **Figures 5B–5D**, t-distributed stochastic neighbor embedding (t-SNE) analysis revealed 9 distinct cellular clusters, comprising epithelial cells, macrophages, mast cells, neutrophils, B cells, dendritic cells, fibroblasts, and endothelial cells. Consistent with the RNA-seq findings, the proportion of macrophages was significantly elevated in the *C*. *albicans*-infected group (**Figure 5E**). Importantly, this increase in macrophages is also observed in human GC samples^22,23^ (**Figures S5C and S5D**). We further divided the macrophage populations into 4 subpopulations based on the Cellmaker2.0 database^24^. Among these subpopulations, we found that cluster 0, which expressed Chil3 and is characterized as M2-like macrophages, was predominantly observed in mice infected with *C*. *albicans* (**Figures 5F–5I**). In contrast, other clusters, including clusters 1, 2, and 3, which expressed H2-Eb1 (M1-like macrophages), were decreased (**Figures 5F–5I**). To validate the macrophage fractions, we employed flow cytometry, using CD206 as a marker for M2-like macrophages and CD80 for M1 macrophages^25^ (gated on Live/Dead^-^, CD45^+^, CD11b^+^, F4/80^+^ markers; validated in **Figure S5E**). We found that the proportion of total macrophages and CD206^+^ M2-like macrophages was significantly elevated in the gastric tumors from *C*. *albicans*-infected, *K19*-*C2mE* mice, while the proportion of CD80^+^ M1-like macrophages showed a contrasting trend (**Figure 5J**), consistent with the results obtained from scRNA-seq analyses. As an additional control, in the gastric lesions collected from germ-free mice infected with *C*. *albicans*, we consistently observed a notable increase in the proportion of macrophages, accompanied by enhanced infiltration of M2-like macrophages and a decreased proportion of M1-like macrophages (**Figure S5F**). Furthermore, in inosine-treated *K19*-*C2mE* mice, we observed a higher proportion of CD206^+^ M2-like macrophages and a lower proportion of CD80^+^ M1-like macrophages in the gastric tumors (**Figure 5K**). Therefore, both *C*. *albicans* and inosine induced M2-like polarization of macrophages during gastric carcinogenesis.

**Figure 5.**
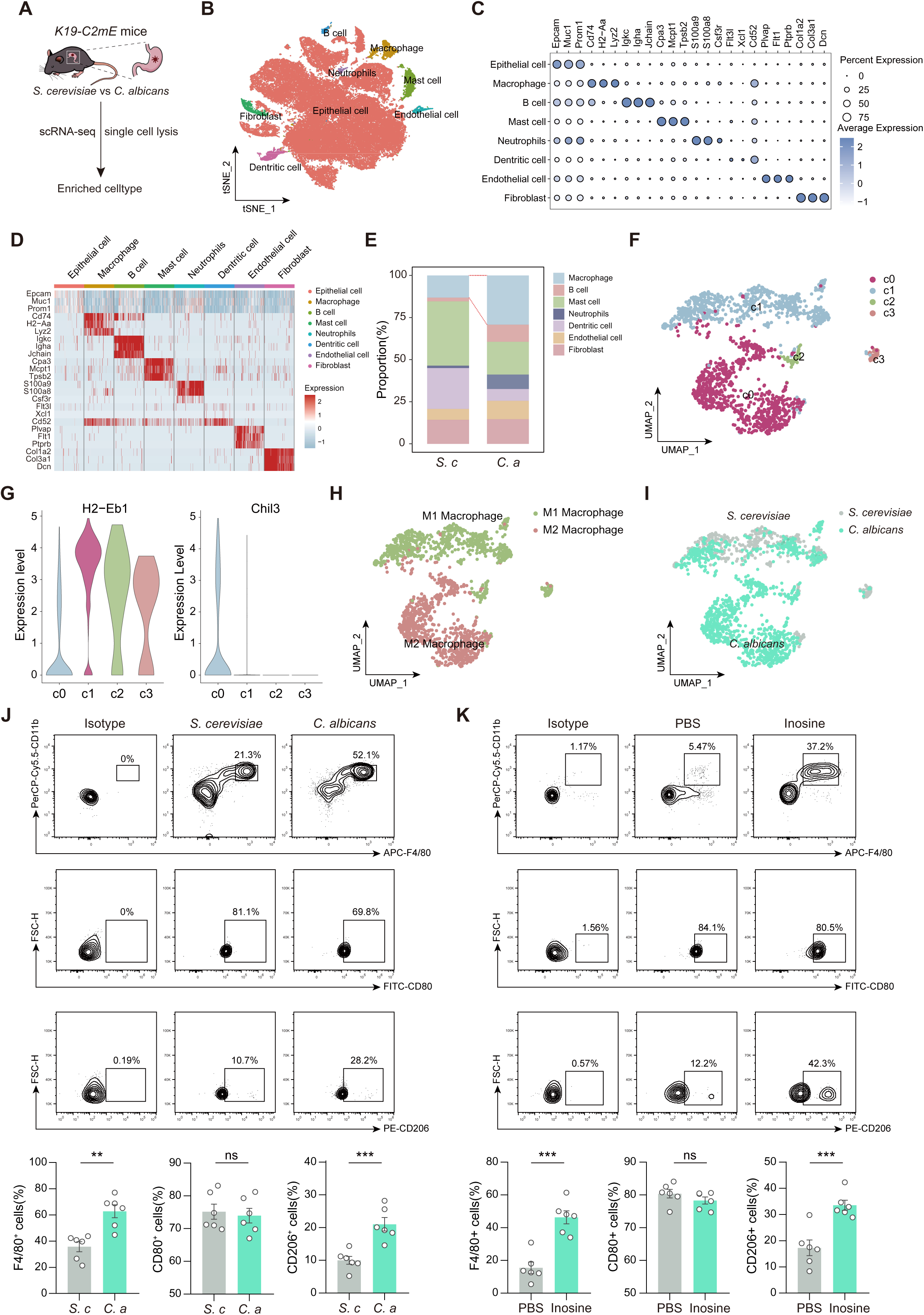
Single-cell sequencing reveals that *C.albicans* mediates protumor through M2 macrophages. (A) Schematic diagram of the procedure for single-cell RNA sequencing of gastric tissue in *S. cerevisiae* or *C. albicans* infected mice (n = 3/group). (B) Visualization of single-cell RNA sequencing data using unsupervised hierarchical clustering and uniform manifold approximation and projection (UMAP), colored by major cell clusters. (C) Dot plot depicting the relative expression of canonical cell type marker genes (annotated per CellMarker 2.0) across major identified clusters. (D) Heatmap showing the relative expression of canonical marker genes (annotated per CellMarker 2.0) across major cell clusters. (E) The relative proportion of immune and stroma cells between *S. cerevisiae* and *C. albicans*-infected *K19-C2mE* mice. Macrophages showed a significant increase in the *C. albicans*-infected group. (F) UMAP plot of macrophages. (G) Violin plot of different gene expression on sub-clusters of macrophages (annotated per CellMarker 2.0) (H-I) Clusters are annotated with predicted cell types according to CellMarker2.0 (G) and distribution of M1 or M2 macrophage across *S. cerevisiae* and *C. albicans*-infected *K19-C2mE* mice (I). (J) Flow cytometry analysis of M1 or M2 macrophage (CD80 or CD206) proportions between *S. cerevisiae* and *C. albicans*-infected *K19-C2mE* mice. (K) Flow cytometry analysis of M1 or M2 macrophage (CD80 or CD163) proportions between PBS or inosine-administered *K19-C2mE* mice. Data were presented as mean ± SEM. Each point represents one subject. Student’s t test (J and K) was used to assess the statistical significance between groups, ns, nonsignificant, ** p < 0.01, *** p < 0.001.

We next investigated the mechanisms through which *C*. *albicans* contributes to the polarization of macrophages toward an M2-like phenotype. Previous research has shown that inosine can regulate the differentiation of regulatory T cells^21,26^; however, its effects on macrophage polarization remain unclear. We therefore used THP-1 cells, RAW 264.7, and bone marrow-derived macrophages (BMDMs), which are cell lines representing macrophage precursors or mature macrophages, and differentiated THP-1 into M0 macrophages *in vitro* by PMA treatment. We found that treating THP-1 and Raw264.7 cells with 100□μM inosine for 5 days, a concentration comparable to that observed *in vivo* after *C. albicans* infection, effectively promoted M2-like polarization, as evidenced by elevated expression levels of CD163 and CD206 (**Figures S6A–S6F**). However, prolonged *in vitro* culture can predispose BMDMs to spontaneous M2 polarization, leading to an increased sensitivity to IL-4^27^ (**Figures S6C and S6F**). We therefore tested a higher concentration to shorten the treatment, and found that treating BMDMs with 1□mM inosine for 24□h effectively induced M2-like polarization, consistent with the effects observed in THP-1 and Raw264.7 cells (**Figures 6A–6C**). Given that 1□mM inosine is within the effective range for the A2A receptor^28,29^, this concentration was used for all subsequent *in vitro* experiments. We found that inosine significantly promoted the polarization of these cells towards the M2-like phenotype, similar to that observed with IL-4 treatment, that are positive control for M2-like polarization (**Figures 6A–6C**). In addition, inosine decreased M1 polarization, as indicated by reduced expression of CD80 and IL-1β, which is in contrast with the effects of LPS and IFN-γ that are canonical inducers of M1 macrophage polarization (**Figures S6G–S6I**). Consistent with the known GC□promoting function of M2 macrophages^30,31^, we observed significantly higher rates of both cell proliferation and migration in GC tumor cells, including AGS, MKN45, and ATK cells, when treated with culture media collected from inosine-treated macrophages (**Figures 6D and 6E, Figures S6J–S6P**).

**Figure 6.**
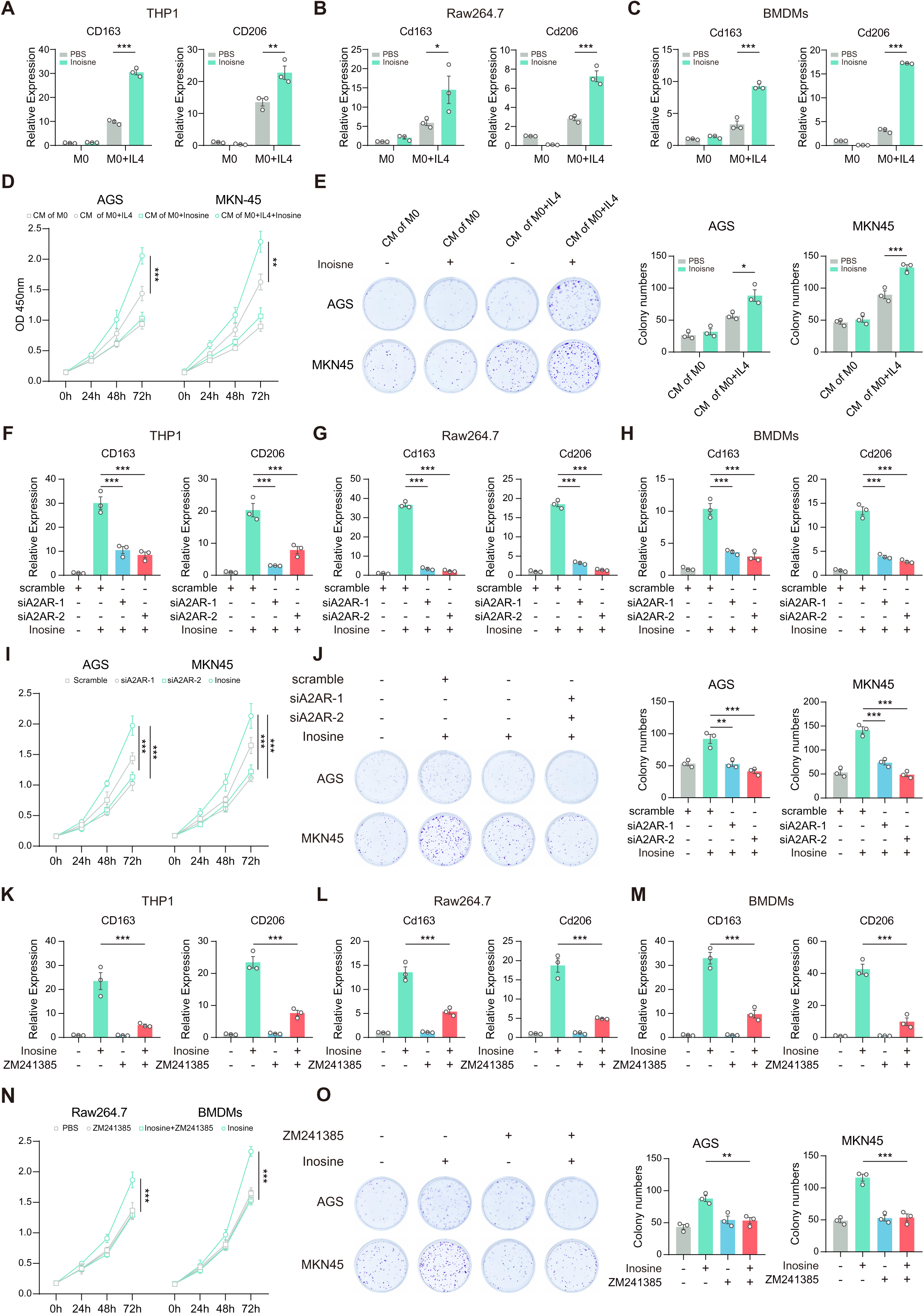
Inosine activates A2AR receptors in macrophages to trigger tumorigenesis. (A-C) RT-qPCR analysis of M2 markers (CD206, CD163) in THP-1 (A), RAW264.7 (B), and bone marrow-derived macrophages (BMDMs) (C) treated with or without inosine (1 mM) for 24 hours under M2-polarizing conditions. (D-E) CCK-8 assay (D) and colony formation assay (E) assessing the effects of conditioned media from differentially polarized-macrophages on tumor cell proliferation. (F-H) RT-qPCR analysis of CD206 and CD163 expression to assess the role of A2aR in inosine-mediated effects. THP-1 (F), RAW264.7 (G), BMDMs (H) were transfected with scramble or A2aR-targeting siRNA (siA2aR) and treated with or without inosine (1 mM) for 24 hours under M2-polarizing conditions. (I-J) CCK-8 assay (I) and colony formation assay (J) assessing the effects of conditioned media from differentially polarized macrophages, with or without A2AR silencing, on tumor cell proliferation. (K-M) RT-qPCR analysis of M2 markers (CD206, CD163) in THP-1 (F), RAW264.7 (G), and BMDMs (H) treated with PBS or ZM241385 (5µM), in the presence or absence of inosine (1 mM) under M2-polarizing conditions. (N-O) CCK-8 assay (N) and colony formation assay (O) assessing the effects of conditioned media from differentially polarized macrophages, with or without ZM241385 (5µM), on tumor cell proliferation. Data were presented as mean ± SEM. Each point represents one subject. One-way ANOVA (A, B, C, D, E, F, G, H, I, J, K, L, M, N, and O) was used to assess the statistical significance between groups, * p < 0.05, ** p < 0.01, *** p < 0.001.

We further investigated the mechanism by which inosine regulates macrophage polarization. Members of the adenosine receptor (AR) family, including A1R, A2aR, A2bR, and A3R^20^, have been identified as receptors for inosine and have previously been shown to regulate tumor formation^32^. We knocked down A2aR, A2bR, and A3R, as well as A1R, which is another family member of AR, in THP-1 cells (**Figure S7A**). As shown in **Figures 6F–6H**, the knockdown of A2aR significantly reduced the enhancing effect of inosine on M2-like macrophage polarization, while the other AR family members did not show this effect (**Figures S7B and S7C**). Mechanistically, A2aR primarily activates the Gαs signaling pathway, increasing cAMP levels and subsequently activating PKA^33,34^ to promote M2 macrophage polarization^34,35^. Consistently, we found that inosine robustly activated the cAMP-PKA pathway (**Figure S7D**). Additionally, we observed that silencing A2aR diminished the capacity of M2 macrophages, through their culture media, to induce proliferation and migration of GC cells (**Figures 6I and 6J, Figures S7E–S7L**). Consistently, the treatment of macrophages with ZM241385, a specific inhibitor of A2aR^36^, significantly inhibited inosine-induced M2-like macrophage polarization (**Figures 6K–6M, Figure S8A**) and decreased these cells’ ability to promote the proliferation and migration of GC cells (**Figures 6N and 6O, Figures S8B–S8I**). These findings indicate that inosine facilitates polarization of macrophages toward an M2-like phenotype through A2aR signaling during gastric tumorigenesis.

### *C. albicans* promotes gastric tumorigenesis through inosine-A2aR signaling in macrophage

We next asked whether *C*. *albicans* contributes to gastric tumorigenesis by facilitating the polarization of M2 macrophages. To test this, we treated *C*. *albicans*-infected *K19*-*C2mE* mice with ZM241385, to block the inosine-A2aR signaling pathway, or we depleted macrophages using clodronate liposomes (Col-lip)^37^ (**Figure S9A**). As expected, both ZM241385 and Col-lip treatment resulted in a significant reduction of M2-like macrophages in the gastric tissue of *K19*-*C2mE* mice following *C*. *albicans* infection (**Figure S9B**). We also observed a marked decrease in tumor count, area, and tumor cell proliferation rates in these treatment groups (**Figures S9C–S9E**), indicating that blocking inosine-A2aR signaling in macrophages impairs the pro-tumor effects mediated by *C*. *albicans*. Furthermore, the well-differentiated adenocarcinoma (WDA) seen in the *C. albicans*-infected mice was no longer present after treatment with either ZM241385 or Col-lip; instead, we observed low-grade dysplasia (**Figure S9B**).

We also examined the tumor□promoting effects of macrophage inosine-A2aR signaling in the inosine-gavaged *K19*-*C2mE* mice (**Figure S9F**). Consistently, we observed a decrease in tumor burden, along with a reduction in dysplastic transformation of gastric mucosa and reduced cell proliferation rates (**Figures S9G–S9J**). Therefore, we demonstrated that *C*. *albicans* elevates inosine levels in the tumor microenvironment, which, in turn, activates the A2aR receptor on macrophages, inducing their polarization toward an M2-like phenotype, and thereby promoting tumor development.

### *C. albicans* and inosine levels resemble GC severity in humans

We examined the levels of *C*. *albicans* and inosine in humans with GC. Serum samples collected from both healthy donors and patients diagnosed with GC showed a significant increase in *C*. *albicans*-specific IgG levels in the serum of GC patients (**Figure 7A; Table S3**) and a seropositivity rate of ∼35% in the GC cohort, compared to ∼5% in healthy donors (**Figure 7B**), suggesting that there is a higher population of *C*. *albicans* in these patients. Supporting this finding, as shown in **Figure 7C**, the ROC curve analysis demonstrated that *C*. *albicans*-specific IgG levels effectively distinguished gastric cancer patients from healthy individuals. These results are consistent with previous ITS sequencing data from GC tissues^15^ and analyses from the TCGA database^16^. Moreover, tumor tissues from seronegative GC patients contained significantly lower levels of fungal DNA than those from seropositive patients (**Figures S10A and S10B**). In addition, we found significantly elevated inosine levels in the peripheral blood of GC patients (**Figure 7D**), with particularly pronounced increases in those infected with *C*. *albicans*, whereas inosine levels in *C. albicans*-negative patients were comparable to those of healthy controls (**Figure 7E**). In particular, there was a strong positive correlation between *C*. *albicans*-specific IgG levels and inosine concentrations (**Figure 7F**) in GC patients. Furthermore, in GC patients who were positive for *C. albicans*, higher abundances of *C. albicans* and inosine were positively associated with more advanced clinicopathological grades (**Figures 7G–7J**) and metastasis status (**Figures 7K–7L, Figures S10C and S10D**), whereas no such associations were observed in *C. albicans*-negative patients (**Figure 7L and Figure S10E**). Consistent with observation in mice and cell lines (**Figures 4K and 4L**), GC patients with higher levels of *C. albicans* colonization exhibited elevated ADA expression in the gastric tissues adjacent to tumors, whereas ADA expression in tumor tissues was significantly lower than in matched non-tumor tissues (**Figures S10F–10H**). Similarly, we observed that increased expression of A2aR correlated with poor prognosis, as evidenced by reduced overall survival, first progression, and post-progression survival (**Figures S10I–10K**). We also discovered that patients were seropositive for both *H*. *pylori* and *C*. *albicans* exhibited higher inosine level worse pathological grades and metastasis compared to those were positive for either *H*. *pylori* or *C*. *albicans* alone (**Figures 7M–7O**). This reinforces the synergistic effects of *H*. *pylori* and *C*. *albicans* in gastric tumorigenesis.

**Figure 7.**
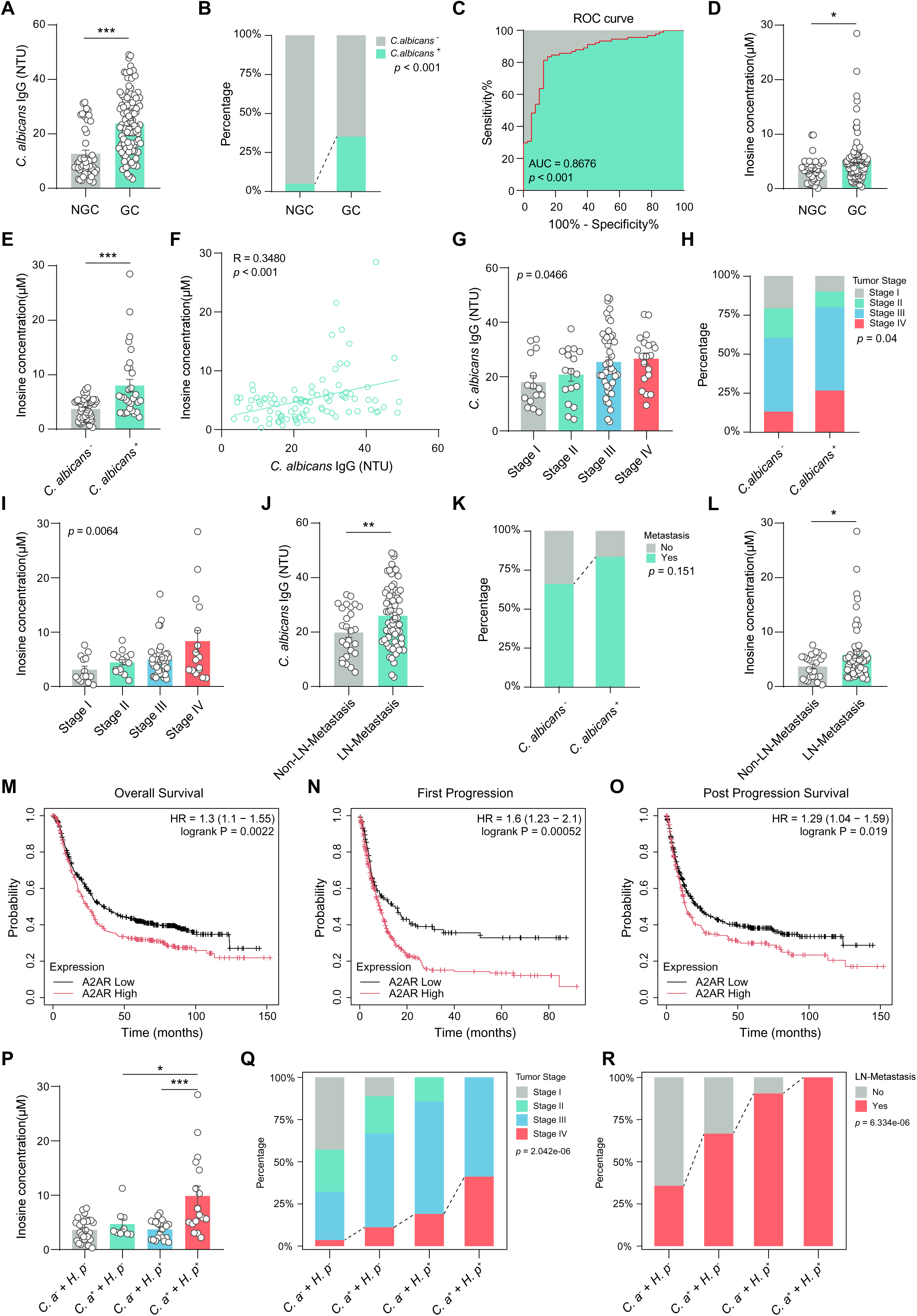
Clinical significance of *C. albicans* levels, and inosine dependence in human GC. (A) Elisa assay of *C. albicans* IgG concentration in serum between non-GC patients (n = 49) and GC patients (n = 103). (B) The proportion of *C. albicans* negative and positive sample between non-GC patients (n = 40) and GC patients (n = 91). (C) The ROC curve for serum anti-*C. albicans* IgG in distinguishing GC (n = 103) from non-GC (n = 49) patients. (D) Elisa assay of inosine concentration in the serum between non-GC patients (n = 38) and GC patients (n = 93). (E) The serum inosine concentration in *C. albicans* negative (n = 54) or positive (n = 30) GC patients. (F) The correlation of the concentration of *C. albicans* IgG and inosine concentration (n = 93) in GC patients. (G) The relationship between the concentration of *C. albicans* IgG and pathological stage in GC patients. (H) The proportion of pathological stage in *C. albicans* negative (n = 55) or positive (n = 30). (I) The relationship between the inosine concentration and pathological stage in GC patients. (J) The relationship between the concentration of *C. albicans* IgG and lymph node metastasis in GC patients. (K) The proportion of lymph node metastasis in *C. albicans* negative (n = 53) or positive (n = 30). (L) The relationship between the inosine concentration and lymph node metastasis in GC patients. (M-O) Overall survival (M), first progression (N), and post-progression survival (O) curves comparing patients with low and high expression of A2AR mRNA in gastric cancer samples from the Kaplan-Meier dataset. (H-I) The inosine level, pathological stage and lymph node metastasis in different *C. albicans* and *H. pylori* infected GC patients. Data were presented as mean ± SEM. Each point represents one subject. Student’s t test (A, D, J, and L), One-way ANOVA (G, I and P), log (rank Mantel-Cox test) (M, N, and O), Kruskal-Wallis rank sum test (Q) and Chi-square test (R) were used to assess the statistical significance between groups, * p < 0.05, ** p < 0.01, *** p < 0.001.

## Discussion

We have identified *C. albicans* as a direct carcinogen in gastric tumorigenesis. In both wildtype mice and *K19*-*C2mE* spontaneous GC mice, we found that *C. albicans* alone is sufficient to induce preneoplastic progression and GC formation. Notably, the pro□tumor efficacy of *C*. *albicans* is comparable to that of the well-established carcinogen *H*. *pylori*, and its clinical prevalence in GC patients (∼35%) is similarly substantial. In addition, co□infection with both pathogens synergistically exacerbates preneoplastic progression and tumor development, reinforcing the notion that gastric carcinogenesis may stem from polymicrobial interactions rather than a single pathogen^4–8^. Our findings could also partially explain the disease heterogeneity observed clinically, especially in GC individuals who do not present with *H*. *pylori* infection.

Of note, compared with bacteria, fungi represent a smaller portion of the total microbiome, with reported read proportions of approximately 10□□ for fungi versus 10□² for bacteria^38^. Specifically, *C. albicans*, an opportunistic commensal fungus, typically constitutes only about 0.1% of the total gastrointestinal microbial abundance in healthy individuals^39–41^. In this context, it plays a role in maintaining gastrointestinal immune balance^42,43^. However, under certain pathological conditions, *C. albicans* can proliferate and increase its abundance, transitioning from a symbiotic organism to a factor that drives disease^39,44^. In the case of GC, we also observed a similar trend: *C. albicans* is the most abundant fungal species, making up about 22% of the total fungal abundance^15^; and it has the highest prevalence in GC tissues, with an estimated occurrence of around 40%^16^. In particular, given its ubiquitous distribution^39,45^, and in this study, we found that the presence of *C*. *albicans* is positively correlated with GC severity and metastasis, suggesting it may serve as a biomarker for predicting GC.

Mechanistically, we show that the pro-GC effects of *C. albicans* depend strictly on its viability and physical interaction with gastric epithelial cells. First, studies have shown that *C. albicans* can adhere to gastric epithelial cells and undergo morphological conversion from yeast to hyphae, which is indicative of its invasive behavior^46,47^. This is consistent with the stable presence of *C. albicans* detected in our GC samples. Second, hypha-associated adhesins, such as Als3 and Hwp1, facilitate the attachment to and invasion of host cells by *C. albicans*^39,46^. These adhesins were also observed in our study of *C. albicans*-infected gastric epithelial tissues. Additionally, the formation of hyphae can mechanically disrupt the epithelium and secrete virulence factors such as candidalysin, leading to epithelial activation and inflammatory responses that may progress to gastritis^48,49^. Importantly, all of these dynamic processes are strictly dependent on the viability of *C. albicans* and cannot be replicated by inactivated *C. albicans*. Furthermore, any method that disrupts the role of the hyphae in adhering to, invading, or activating epithelial cells—such as heat inactivation of *C. albicans*, eradication of *C. albicans* using the antibiotic amphotericin B (AmB), or separating *C. albicans* from the epithelial cells—will reduce its pro-GC effects.

Addictionlly, we identified inosine as a key metabolic mediator linking *C. albicans* colonization to the establishment of an immunosuppressive tumor microenvironment. *C. albicans* selectively triggers inosine production in normal gastric mucosal epithelial cells (by elevating ADA expression) after colonization, but not in established GC cells. This spatiotemporal pattern implies that inosine accumulation is an early epithelial event preceding malignant transformation. Downstream of this production, inosine drives immunosuppression by polarizing macrophages toward an M2-like phenotype, which is recognized as a prerequisite for GC development^50^. We further demonstrate that inosine drives immunosuppression by polarizing macrophages toward an M2□like phenotype. It has previously been shown that M2□like macrophages play a crucial role in establishing an immunosuppressive environment in GC: they produce anti□inflammatory cytokines^51,52^ (e.g., IL□10, TGF□β) and express immune□checkpoint ligands, thereby impairing CD8□ T□cell infiltration and function while recruiting other immunosuppressive cells such as regulatory T cells and myeloid□derived suppressor cells^53,54^. We found that it is the activation of A2aR, a type of purinergic receptor in macrophages, by inosine, that promotes their M2 polarization. Importantly, inhibition of A2aR signaling can reverse the M2 polarization and the immunosuppressive microenvironment caused by both *C*. *albicans* and inosine, and suppress their tumorigenic effects. Interestingly, our findings underlie the context□dependent function of inosine, and may explain its dual role in tumor immunology in other types of tumor^20,21,26,32^. For example, it was shown that in immunocompetent models (e.g., during CAR□T therapy), inosine can enhance Th1 responses and sustain T□cell stemness^20,21,55^. This divergence likely reflects differences in timing, microenvironmental cues, and immune□cell composition. In GC, *C*. *albicans*□induced inosine accumulation occurs early, favoring the establishment of an immune□excluded tumor microenvironment. Furthermore, we acknowledge that elevated inosine levels could arise through mechanisms unrelated to *C. albicans*. In particular, previous studies have shown that elevated inosine levels may arise through mechanisms unrelated to *C. albicans*. Certain mechanisms may involve specific bacterial species, such as *Bifidobacterium pseudolongum*, which can produce inosine^21^. Additionally, host cellular stress responses, such as the release of inosine from brown adipocytes undergoing apoptosis, or dysregulated purine metabolism in tumor cells—particularly in the breast, colon, kidney, liver, and stomach^21,32,56,57^—can also play a role. Such pathways could also potentially contribute to GC progression in other circumstances.

Beyond its role in initiating early inosine production from normal epithelium, we also found *C. albicans* residing as an active intratumor mycobiome in both *K19-C2mE* mice and human GC tissues. In other types of cancer, for instance, some pan-cancer analyses have shown that while the intratumor mycobiome accounts for relatively low abundance (approximately 4–13% of the total multi-kingdom microbiome, similar to that of *C. albicans*^16,58^, it can exert significant biological effects in tumorigenesis. For example, *Malassezia* has been shown to promote pancreatic cancer progression via the activation of the mannose-binding lectin pathway^59^. *Alternaria alternata* induces tumor cells to secrete IL-33, recruiting TH2 and ILC2 cells to establish an immunosuppressive microenvironment^60^. Similarly, *Aspergillus sydowii* enhances tumor growth through interactions with myeloid-derived suppressor cells and T cells in lung cancer^61^. These findings support that the intratumor mycobiome can play a significant role in cancer development. Thus, *C. albicans* may also exert distinct, localized tumor-promoting effects within GC tissues.

The pathogenicity of *C. albicans* is also profoundly influenced by the broader gastric microenvironment, particularly pH and the bacterial microbiome. Elevated pH levels have been shown to promote hyphal transition, enhance adhesion, and facilitate mucosal invasion^39,46,47^, thereby amplifying the pro-GC effects of *C. albicans* compared to more acidic conditions in a healthy stomach, where *C. albicans* predominantly exists as a commensal yeast with limited virulence. Moreover, broad-spectrum antibiotics, such as those used for *H. pylori* eradication, can disrupt protective commensal bacteria, such as *Lactobacillus*, which normally help maintain an acidic environment and suppress fungal growth. The loss of these microbial constraints may further facilitate overgrowth of *C. albicans* and increase its pathogenicity^62–64^. In addition to gastric tumorigenesis and antibiotic exposure, *C. albicans* can actively alkalinize its local environment by producing ammonia, further supporting its colonization^65,66^. We confirmed this alkalinizing effect in *C. albicans*-infected *K19-C2mE* mice, aged C57BL/6 mice, and germ-free mice. Importantly, regarding clinical settings, gastric pH is often elevated in patients with gastric cancer due to factors such as long-term antibiotic use, proton pump inhibitors, or *H. pylori* eradication therapy. Therefore, in clinical practice, focusing solely on gastric acid suppression or *H. pylori* eradication while neglecting to address fungal dysbiosis (e.g., involving *C. albicans*) may be inadequate for interrupting gastric carcinogenesis. In high-risk populations for GC—particularly those with prolonged use of acid-suppressive agents or antibiotics—monitoring and regulating gastric fungal colonization (especially that of *C. albicans*) could be critically important.

Although our data suggest that *C. albicans*, without causing significant changes in gastric bacterial composition, can promote gastric tumorigenesis in the *K19-C2mE* mouse model, we cannot rule out the possibility that bacteria may play a role in gastric tumorigenesis in patients. First of all, the *K19-C2mE* mice used were housed under specific pathogen-free (SPF) conditions, limiting their exposure to bacterial pathogens. This is quite different from GC patients, who encounter various pathogens. For example, it was shown that *K19-C2mE* mice^19^ have intrinsically low levels of gastric bacteria, averaging around 10□ CFU per stomach. This amount is significantly lower than what is found in distal gastrointestinal regions, such as the colon, where bacterial densities can reach 10¹¹-10¹² CFU/mL. It was later suggested that the highly acidic stomach environment (pH < 4) restricts the bacterial community to a limited number of acid-tolerant taxa, including *Lactobacillus*^67^, *Streptococcus*^4^, and *H. pylori*^68^. Therefore, the intrinsically low abundance of bacteria and, consequently, low diversity within bacterial communities may limit the potential for community remodeling following *C. albicans* infection. In comparison, GC patients, particularly those with atrophic gastritis, may experience an alkalinization of the gastric environment^69^. This change could alter bacterial communities prior to the development of GC, potentially allowing bacteria to contribute to the formation of cancer afterwards^4,13^.

Our findings also carry direct translational implications. First, the high prevalence and stage□correlation of *C*. *albicans* suggest its utility as a non□invasive biomarker for risk stratification and early detection. In particular, detection of *C*. *albicans*, and possibly *H*. *pylori*, may assist in better distinguishing lymph node metastasis in patients with GC, because traditional contrast-enhanced computed tomography (CT) scanning or magnetic resonance imaging (MRI) for lymph node enlargement can be misleading, as it may be confounded by unrelated factors like bacterial infections. Identifying these biomarkers may help guide treatment decisions, including whether to perform lymph node dissection or initiate neoadjuvant therapy. Second, given that GC is frequently resistant to immunotherapy due to the immunosuppressive microenvironment, eradicating *C*. *albicans* (e.g., via antifungal agents) could sensitize the immunotherapy. Third, targeting the inosine□A2aR axis-for instance, with A2aR antagonists already under clinical investigation in other cancers-may offer a rational immunometabolic strategy to reverse macrophage□mediated immunosuppression and enhance the efficacy of immunotherapeutic approaches. Finally, as dysregulated purine metabolism (including inosine) has also been linked to chemotherapy resistance^70^, adjuvant modulation of this pathway may improve responses to conventional agents such as fluorouracil and platinum.

In summary, our work establishes *C*. *albicans* as a clinically prevalent, direct gastric carcinogen and delineates a fungal□metabolic□immune axis that drives tumorigenesis. These insights expand the etiological framework of GC and reveal novel opportunities for microbiome□guided prevention, risk stratification, and immunometabolic therapies for this type of cancer.

### Limitations of the study

Several limitations of this study should be acknowledged. First, although we delineate the crosstalk between C. albicans and macrophages within the gastric cancer microenvironment, potential direct interactions between C. albicans and tumor cells were not investigated and remain to be addressed in future work. Second, the precise mechanisms by which C. albicans colonizes gastric epithelial cells remain poorly defined and warrant further mechanistic dissection. Third, while we detected C. albicans within gastric tumor tissues of both K19-C2mE mice and patient samples, the specific role of this intratumor fungal population in gastric carcinogenesis remains unclear. Despite these limitations, our study provides evidence that fungal members of the gastric microbiome can actively contribute to gastric tumorigenesis.

## ACKNOWLEDGMENTS

We thank the Center of Clinical Laboratory at Zhongshan Hospital of Xiamen University for assistance with the identification and culture of clinically derived *Candida albicans*. We thank Dr. Hiroko Oshima for kindly providing the *K19-C2mE* transgenic mice. We thank Lei Zhang, School of Life Sciences, Core Facility of Biomedical Sciences, Xiamen University, for the kind support in untargeted metabolomics. This work was financially supported by the National Natural Science Foundation of China (82122046 to X.H., 82103160 to J.Z., 82500246 to Z.M.), the Young Top Talents of Fujian Young Eagle Project, the Fujian Health Youth Scientific Research Project (2021ZQNZD018).

## AUTHOR CONTRIBUTIONS

J.T. and Y.X. conducted the experiments and drafted the manuscript. A.C assisted with mouse experiments. Z.L. performed histological evaluation as a pathologist. J.T. and M.Z. conducted bioinformatics analyses. G.P. assisted with flow cytometry. X.H., Y.Z. and W.W. provided clinical samples from gastric cancer patients. S.Z. assisted with the *Candida* IgG and inosine detection. J.Z., J.M., and H.Z. provided constructive suggestions and support. X.H. and G.S. conceived the project, formulated hypotheses, and designed the studies. X.H., and C. Z. reviewed and revised the manuscript.

## STAR★METHODS

### KEY RESOURCES TABLE

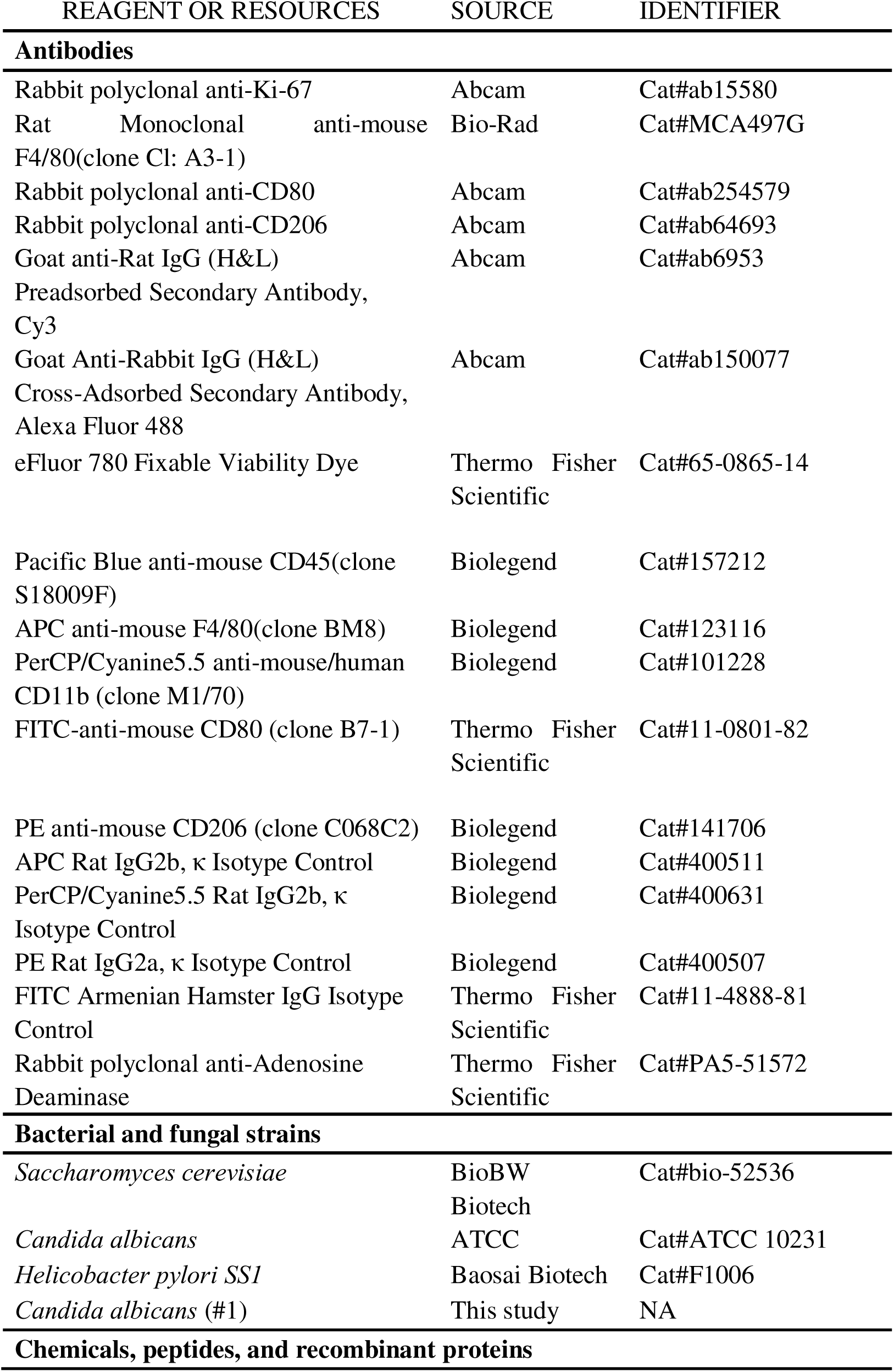
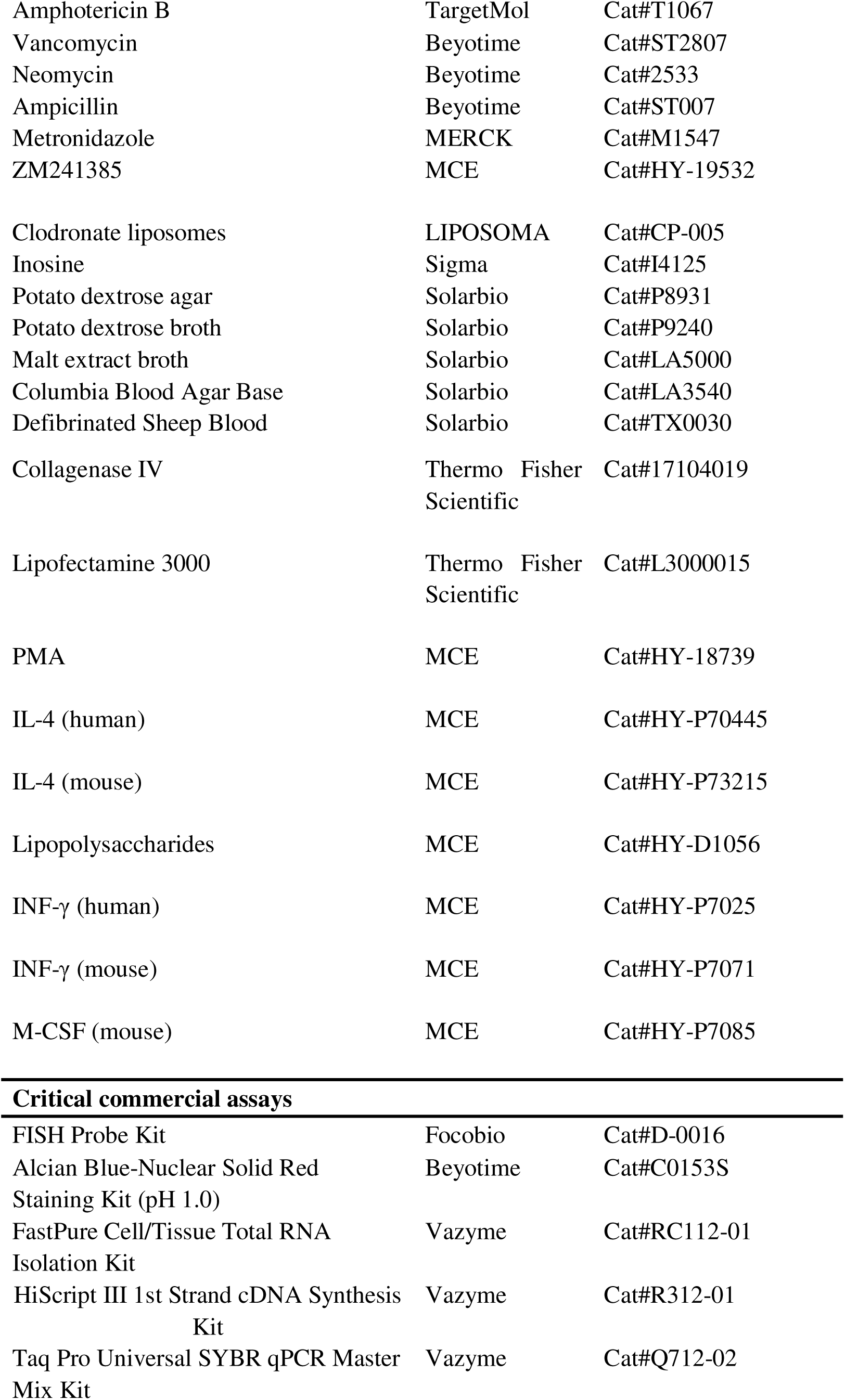
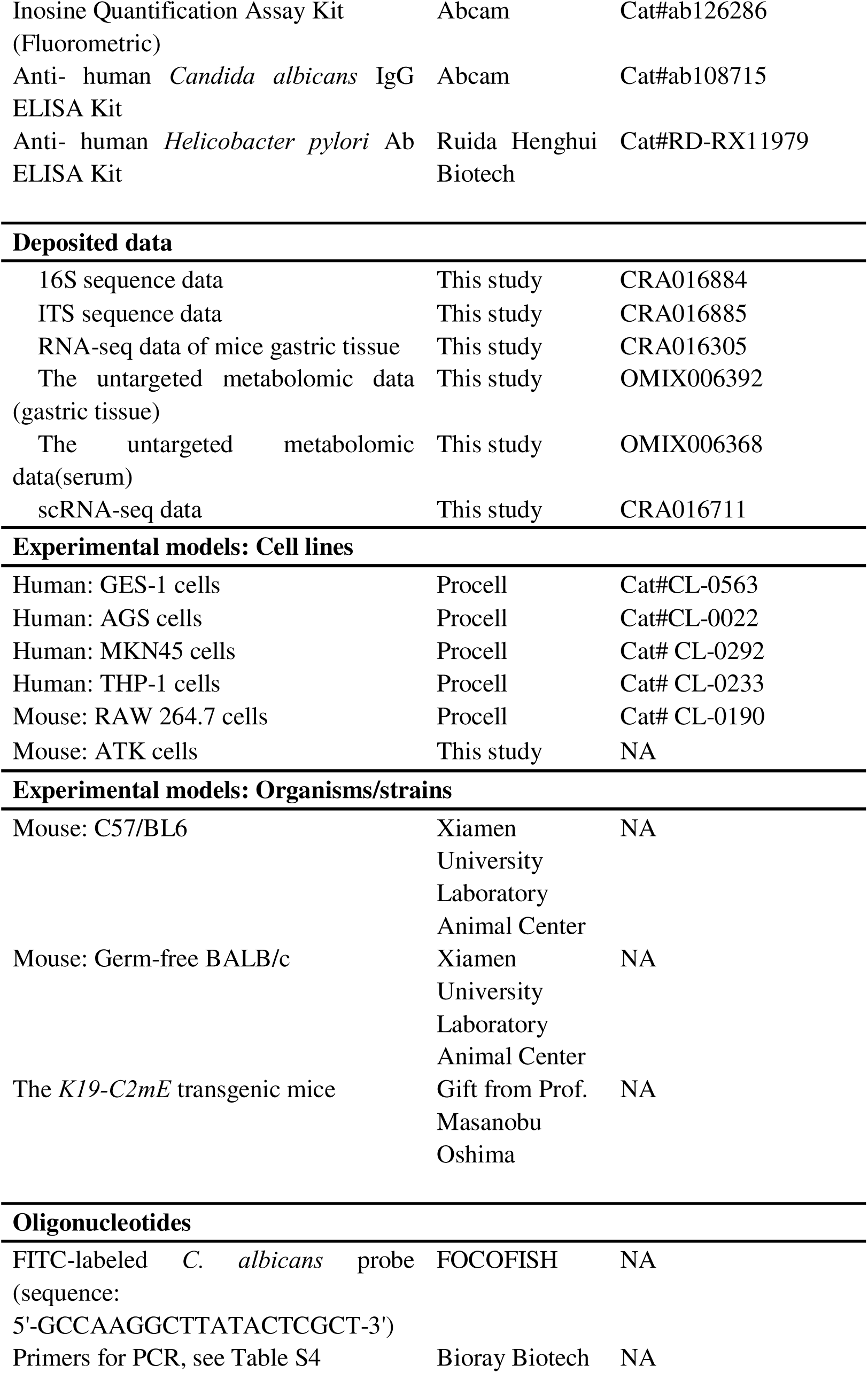
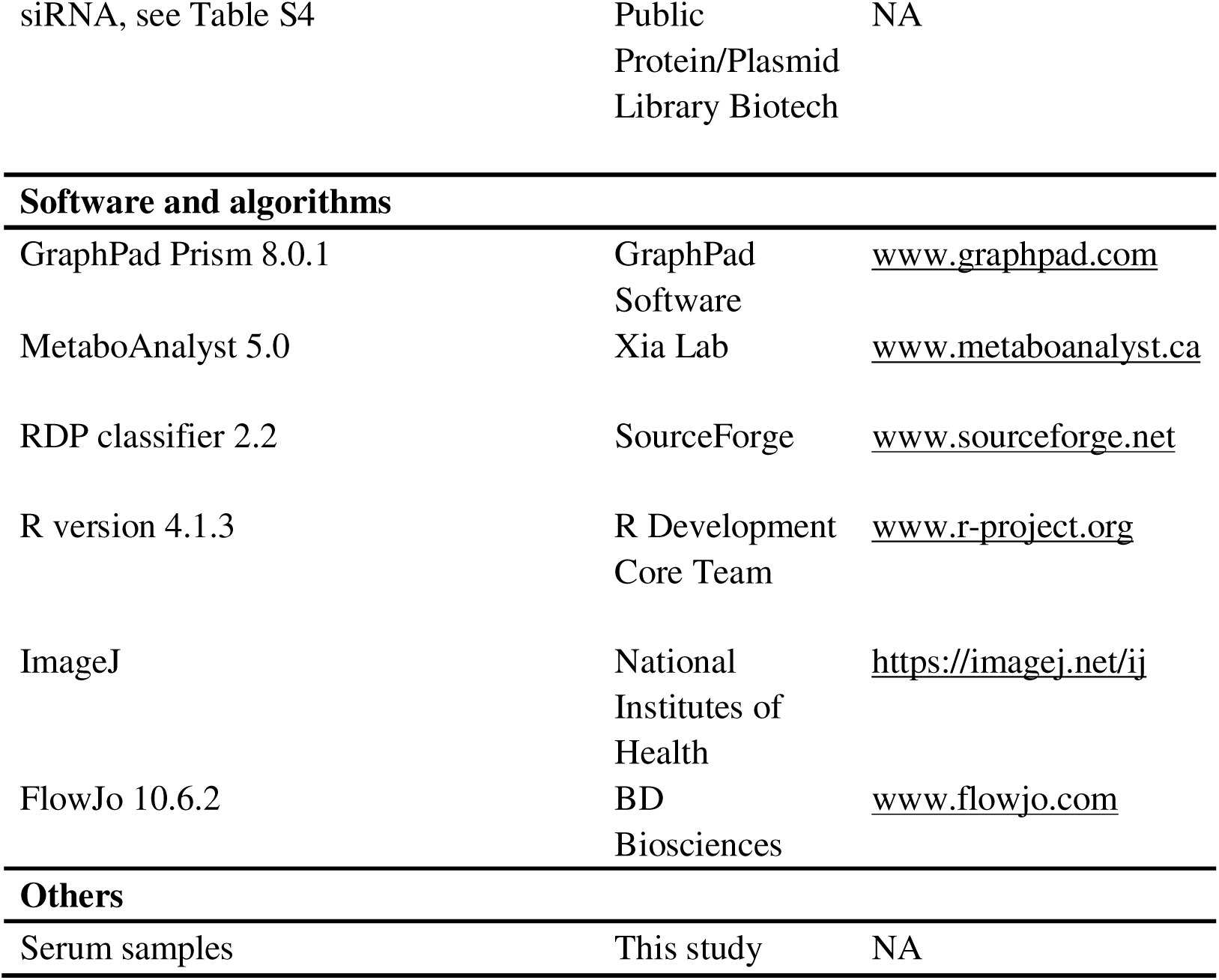

### RESOURCE AVAILABILITY

#### Lead contact

For further information and experimental details, please contact the corresponding author, Dr. Xuehui Hong.

#### Materials availability

This study did not generate new unique reagents.

#### Data and code availability

The raw sequence data reported in this paper have been archived in the Genome Sequence Archive in National Genomics Data Center, China National Center for Bioinformation & Beijing Institute of Genomics, Chinese Academy of Sciences^71,72^. The data includes: Single-cell RNA-seq data of *S. cerevisiae* or *C. albicans*-infected *K19-C2mE* mice gastric tissue: CRA016711; RNA-seq data of *S. cerevisiae* or *C. albicans*-infected *K19-C2mE* mice gastric tissue: CRA016305; 16S sequence data and ITS sequence data of *S. cerevisiae* or *C. albicans*-infected *K19-C2mE* mice gastric tissue: CRA016884 and CRA016885. These datasets are publicly accessible at https://ngdc.cncb.ac.cn/gsa.

The metabolomic data detailed in this paper have been submitted in the OMIX database at the China National Center for Bioinformation & Beijing Institute of Genomics, Chinese Academy of Sciences (https://ngdc.cncb.ac.cn/omix) ^72,73^. The data includes: The untargeted metabolomic data of *S. cerevisiae* or *C. albicans*-infected *K19-C2mE* mice gastric tissue and serum: OMIX006392 and OMIX006368. All data can be obtained from the corresponding authors upon request.

This paper does not report original code.

### EXPERIMENTAL MODEL AND STUDY PARTICIPANT DETAILS

#### Mouse models

##### *C. albicans* infection in C57BL/6 mice

Male C57BL/6 specific pathogen-free (SPF) mice (6-8 weeks old) were randomly assigned to four experimental groups: potato dextrose broth (PDB; negative control), *H. pylori SS1* (positive control), *C. albicans*, and *C. albicans* plus *H. pylori SS1*(co-infection). Mice in the PDB or *C. albicans* groups received oral gavage of broth medium (100 μL) or *C. albicans* (1×10^8^, colony-forming units [CFU]^18^ suspended in 100 μL PDB), respectively, once every three days throughout the infection period. For *H. pylori SS1* infection, mice were orally gavaged with *H. pylori SS1* (2×10^9^ CFU^4^ in 100 PDB) five times over a two-week period, a protocol previously shown to achieve stable gastric colonization^4^. In the fungal-bacterial co-infection model, mice received *C. albicans* and *H. pylori SS1* concomitantly starting from the first inoculation, followed by the same dosing schedules as used in the respective mono-infection groups. Mice from each experimental group were euthanized at 3, 6, 9, and 12 months post-infection for subsequent analyses. All animal procedures were approved by the Animal Experimentation Ethics Committee of Xiamen University (XMULAC20200080) and were performed in accordance with institutional and national guidelines for laboratory animal care.

##### *C. albicans* infection in Germ-free mice

Germ-free BALB/C male mice (6 weeks-old) were randomly assigned to PDB control and *C. albicans*-infected groups. Mice in the PDB or *C. albicans* groups received oral gavage of broth medium (100 μL) or *C. albicans* (1×10^8^, CFU suspended in 100 μL PDB), respectively, once every three days throughout the 9 months infection period. All animal procedures were approved by the Animal Experimentation Ethics Committee of Xiamen University (XMULAC20200080) and were performed in accordance with institutional and national guidelines for laboratory animal care.

##### *K19-C2mE* transgenic mice

*K19-C2mE* transgenic mice, a well-established model of gastric intraepithelial neoplasia, were provided by Prof. Masanobu Oshima^74^. In this model, increased epithelial proliferation is restricted to the glandular stomach (corpus), and tumors originate from this same proliferative epithelium. Tumor areas are marked by red circles and hyperplastic/proliferative regions by white circles. All animal procedures were approved by the Animal Experimentation Ethics Committee of Xiamen University (XMULAC20200080) and were performed in accordance with institutional and national guidelines for laboratory animal care.

For *C. albicans* infection in *K19-C2mE* mice model, *K19-C2mE* mice (15 weeks-old) were randomly assigned to five experimental groups: potato dextrose broth (PDB; negative control), *S. cerevisiae* (negative control), *H. pylori SS1* (positive control), *C. albicans*, and *C. albicans* plus *H. pylori SS1*(co-infection). Mice in the PDB, *S. cerevisiae* or *C. albicans* groups received oral gavage of broth medium (100 μL), *S. cerevisiae* (1×10^8^ CFU^18^ suspended in 100 μL PDB) or *C. albicans* (1×10^8^ CFU suspended in 100 μL PDB), respectively, once every three days throughout the 2-month experimental period. For *H. pylori SS1* infection, mice were orally gavaged with *H. pylori SS1* (2×10^9^ CFU in 100 PDB) five times over a two-week period. In the fungal-bacterial co-infection model, mice received *C. albicans* and *H. pylori SS1* concomitantly starting from the first inoculation, followed by the same dosing schedules as used in the respective mono-infection groups.

For clinical isolate-derived *C. albicans* infection in *K19-C2mE* mice model, *K19-C2mE* mice (15 weeks-old) were randomly assigned to three groups: *S. cerevisiae*, *C. albicans*, and a clinical isolate strain (*C. albicans* #1). Mice in the *S. cerevisiae*, *C. albicans* or *C. albicans*#1 groups received oral gavage of *S. cerevisiae* (1×10^8^ CFU suspended in 100 μL PDB), *C. albicans* (1×10^8^ CFU suspended in 100 μL PDB) or *C. albicans* #1(1×10^8^ CFU suspended in 100 μL PDB), respectively, once every three days throughout the 2-month experimental period.

To determine fungal dependency and microbial specificity, *K19-C2mE* mice (15 weeks-old) were randomly assigned to five experimental groups: *S. cerevisiae* (negative control), *C. albicans*, *C. albicans* plus amphotericin B, *C. albicans* plus antibiotic mixture, and Heat-killed *C. albicans* (H.K *C. albicans*). For fungal clearance, mice received amphotericin B (200 μg per mouse, orally, daily) for five consecutive days, followed by amphotericin B (0.5 μg/mL) administered in drinking water for an additional 7 weeks until study termination^59^. For bacterial depletion, mice were treated with an antibiotic mixture consisting of vancomycin, neomycin, ampicillin, and metronidazole for one week prior to fungal gavage to eliminate bacterial communities^75^. H.K *C. albicans* (1×10^8^ CFU in 100 μL PDB) was prepared by incubating fungal suspensions at 95 °C for 2 hours before gavage. Following these interventions, mice were orally gavaged with either *S. cerevisiae* (1×10^8^ CFU in 100 μL PDB) or *C. albicans* (1 × 10^8^ CFU in 100 μL PDB) according to group assignment throughout the 2-month experimental period.

To assess metabolites specificity, *K19-C2mE* mice (15 weeks-old) were randomly assigned to four experimental groups receiving supernatants derived from: *S. cerevisiae* monoculture, *C. albicans* monoculture, GES-1 gastric epithelial cells co-culture with *S. cerevisiae*, and GES-1 cells co-culture with *C. albicans*. Mice were continuously administered the indicated supernatants as drinking fluids throughout the 2-month experimental period.

For inosine supplementation experiments, *K19-C2mE* mice (15 weeks-old) were randomly assigned to PBS or inosine-treated groups. Mice received daily oral gavage of 100μL PBS or inosine (300 mg/kg^21^ in 100μL PBS) throughout the 2-month experimental period.

To assess the roles of A2A receptor signaling and macrophages in *C. albicans*-driven gastric pathology, *K19-C2mE* mice (15 weeks-old) were randomly assigned to four experimental groups: *S. cerevisiae* (negative control), *C. albicans*, *C. albicans* plus A2AR inhibitor (ZM241385), and *C. albicans* plus macrophage clearance (Clodronate Liposomes). Mice in the A2A receptor inhibition group received intraperitoneal injections of ZM241385 (0.4 μg per mouse, twice weekly). Macrophage depletion was achieved by intravenous administration of clodronate liposomes (200 μL per mouse, three times weekly) Following these treatments, mice were orally gavaged with either *S. cerevisiae* (1 × 10^8^ CFU in 100 μL PDB) or *C. albicans* (1 × 10^8^ CFU in 100 μL PDB) throughout the 2-month experimental period.

To evaluate the involvement of A2A receptor signaling and macrophages in inosine-driven gastric pathology, *K19-C2mE* mice (15 weeks-old) were randomly assigned to four experimental groups: PBS (negative control), inosine, inosine plus A2AR inhibitor (ZM241385), and inosine plus macrophage clearance (Clodronate Liposomes). Mice in the A2A receptor inhibition group received intraperitoneal injections of ZM241385 (0.4 μg per mouse, twice weekly). Macrophage depletion was achieved by intravenous administration of clodronate liposomes (200 μL per mouse, three times weekly) Following these treatments, mice were daily orally gavaged with 100μL PBS or inosine (300 mg/kg in 100μL PBS) throughout the 2-month experimental period.

##### Subject, bacteria, and cell line details

###### Human subjects

Serum samples were collected at the Affiliated Zhongshan Hospital of Xiamen University, including samples from 49 individuals with non-malignant tumors serving as non–gastric cancer controls and 103 patients with gastric cancer. All study procedures involving human participants were reviewed and approved by the Ethics Committee of the Affiliated Zhongshan Hospital of Xiamen University (xmzsyyky 2022-207). Written informed consent was obtained from all participants prior to serum collection. Detailed clinical characteristics of patients and controls are provided in **Table S3**.

###### Fungi and bacteria

*S. cerevisiae* (bio-52536) was obtained from BioBW Biotechnology and cultured in malt extract broth (LA5000; Solarbio) medium at 28 □ with shaking at 160 rpm under aerobic conditions overnight. *C. albicans* (ATCC10231) was obtained from American Type Culture Collection and cultured on potato dextrose agar (P8931, Solarbio). For liquid culture, *C. albicans* was grown in potato dextrose broth (P9240, Solarbio) medium at 28□ with shaking at 160 rpm under aerobic conditions overnight. *H. pylori SS1* was cultured on Columbia blood agar base (LA3540, Solarbio) supplemented with 10% defibrinated sheep blood (TX0030, Solarbio) and incubated at 37 °C for 48 h under microaerophilic conditions using an anaerobic jar.

##### Cell lines

Human gastric cell lines GES-1, AGS, MKN45, and THP-1, as well as murine macrophage Raw264.7 cells, were obtained from Procell Life Science Technology. AGS were cultured in Ham’s F-12 medium (PM150810, Procell). GES-1, MKN45, and THP-1 were cultured in RPMI 1640 medium (PM150110, Procell). Raw264.7 were cultured in DMEM medium (PM150210, Procell). All media were supplemented with 10% fetal bovine serum (10099141C, Thermo Fisher Scientific) and 1% penicillin-streptomycin (S110JV, BasalMedia). Bone marrow-derived macrophages (BMDMs) were generated by isolating bone marrow cells from the femurs and tibias of 6–8-week-old C57BL/6 mice. Cells were filtered and cultured in DMEM (PM150210, Procell) supplemented with 10% FBS and 20 ng/mL M-CSF (HY-P7085, MCE). Adherent BMDMs were used for experiments on day 7 and confirmed by F4/80□CD11b□ staining^76^. ATK cells were isolated from Cldn18^Cre^Apc^fl/fl^Tp53^fl/fl^Kras^G12D^ mice^77^ and cultured in DMEM medium with 10% FBS and 1% penicillin/streptomycin. All cells were cultured at 37°C in a humidified incubator with 5% CO_2_.

### METHODS DETAILS

#### Histological analysis

Fresh gastric tissues were collected, washed with PBS (pH 7.4), and fixed in 4% paraformaldehyde for 24 h. Fixed tissues were dehydrated through a graded ethanol series, embedded in paraffin, and sectioned at 4 µm thickness.

For Hematoxylin-Eosin (H&E) Staining, paraffin sections were deparaffinized, rehydrated through graded ethanol, and stained with hematoxylin. Differentiation was performed in 1% hydrochloric acid alcohol, followed by bluing in LiCO□ solution. Sections were counterstained with eosin, dehydrated, cleared, and mounted with neutral resin prior to imaging. All histopathological evaluations were independently assessed and diagnosed by experienced pathologists in a blinded manner.

For immunohistochemistry (IHC) Staining, sections were deparaffinized, rehydrated, and subjected to antigen retrieval using enzyme-based solution (DIG-3009, MXB Biotech). Immunostaining was performed using the UltraSensitive SP Kit (KIT-9730, MXB Biotech) according to the manufacturer’s instructions. Primary antibody against Ki67 (ab15580, Abcam) was applied overnight at 4 °C. Signals were developed using DAB chromogenic solution (DAB-0031, MXB Biotech). Sections were counterstained with hematoxylin, dehydrated, cleared, and mounted with neutral resin before imaging.

For Alcian Blue/Nuclear Fast Red Staining, paraffin sections were deparaffinized and rehydrated through graded ethanol. Samples were stained with 1% Alcian Blue (pH 1.0) for 30 min, followed by counterstaining with 0.1% Nuclear Fast Red for 10 min (C0153S, Beyotime). After staining, sections were dehydrated, mounted, and imaged according to standard protocols.

#### Fluorescence in Situ Hybridization (FISH)

*C. albicans* colonization was detected using a FISH probe labeled with FAM (sequence: 5’-GCCAAGGCTTATACTCGCT-3’). Fresh murine gastric tissues were collected, rinsed with PBS (pH 7.4), and fixed in 4% paraformaldehyde for 24 h. Fixed tissues were dehydrated, embedded in paraffin, and sectioned at 4 µm thickness. Paraffin sections were deparaffinized, rehydrated, and processed using the FISH Probe Kit (D-0016, Focobio) according to the manufacturer’s instructions. Briefly, sections were pre-incubated in blocking buffer at 55 °C for 2 h. The FAM-labeled *C. albicans* probe, prepared as a 1:50 dilution in 25% hybridization buffer and preheated at 88 °C for 3 min, was applied to the sections and incubated in the dark at 42 °C for 24 h. After washing and dehydration, sections were mounted with DAPI-containing anti-fade solution (P0131; Beyotime). Fluorescence images were acquired using a fluorescent microscope (Leica).

#### Isolation of live fungi from mouse stomach

Fresh murine gastric tissues were homogenized in 50% potato dextrose broth (PDB)-glycerol. Serial dilutions (1:5, 1:25, and 1:50) of the homogenate were plated on sterile potato dextrose agar (PDA) supplemented with antibiotics to suppress bacterial growth. Individual fungal colonies were picked and subjected to MALDI-TOF mass spectrometry (MS) for species identification (Table S1) and further verified by colony PCR to confirm the presence of viable *C. albicans*.

#### Conditioned medium preparation

For fungal conditioned medium, *S. cerevisiae* or *C. albicans* were cultured in RPMI 1640 medium supplemented with 10% FBS at 37 °C for 24 h. The supernatants were collected, filtered through a 0.22 μm sterile syringe filter, and mixed with fresh medium at a 1:1 ratio^78^.

For fungal-epithelial co-culture conditioned medium, *S. cerevisiae* or *C. albicans* were co-cultured with GES-1, AGS, or MKN45 cells in RPMI 1640 medium supplemented with 10% FBS at 37 °C for 24 h. The conditioned media were then collected and filtered through a 0.22 μm sterile syringe filter.

For macrophage conditioned medium, THP-1 cells were differentiated with PMA(100 ng/mL) for 24 h and polarized to M1 macrophages with LPS (100 ng/mL) plus IFN-γ (20 ng/mL) for 24 h, or to M2 macrophages with IL-4 (20 ng/mL) for 24 h. Raw264.7 and BMDMs were similarly polarized to M1 or M2 macrophages under the same conditions. Treatments were applied as indicated, including inosine (pretreated, 1mM), ZM241385(pretreated, 5μM), or siAR for 24 h. Conditioned media were collected, filtered through a 0.22 μm sterile syringe filter, and mixed with fresh medium at a 1:1 ratio.

#### CCK-8 proliferation assay

S. cerevisiae or C. albicans were cultured in their respective media, centrifuged at 1000 rpm for 1 minute after 24 hours, counted, and resuspended in PBS buffer. Equal numbers of S. cerevisiae or C. albicans were cultured in 1640 medium with 10% FBS at 37°C in a humidified cell incubator. After the designated incubation period, fresh conditioned media were obtained using a sterile syringe and a 0.22 μm filter, and mixed with fresh media at a 1:1 ratio to to avoid effects of nutrient/metabolite exhaustion.

AGS, MKN45 and ATK cells were counted, resuspended in the respective conditioned media (1×10³ cells per well), and seeded into 96-well plates. Cell proliferation was measured at 24, 48, and 72 h using a CCK-8 assay kit (HY-K0301; MCE) according to the manufacturer’s instructions.

#### Colony Formation Assay

Fungal broth or fungi at a MOI of 10 were counted and seeded in the upper layer of the co-culture system, while a designated number of AGS/MKN45 cells were seeded in the lower layer. The fungal broth and culture medium were replaced every two days. After 7 days, the cells were fixed with 4% paraformaldehyde (Solarbio, Cat No: P1110) for 30 minutes and stained with 0.1% crystal violet solution (Solarbio, Cat No: G1064) for 15 minutes. The cells were then washed three times with PBS, air-dried, and photographed for statistical analysis. For metabolite treatments, 1% HEPES (1M) (Gibco, Cat No: iCell-01200) was added to the system to maintain stability.

AGS, MKN45, and ATK cells were counted and resuspended in the respective conditioned media. Cells (1×10³ per well) were seeded into 6-well plates and cultured for 14 days to allow colony formation. The medium was refreshed every 3 days. At the endpoint, colonies were fixed with 4% paraformaldehyde for 15 min, stained with 0.5% crystal violet for 20 min, washed with water, and air-dried. Colonies were counted under a light microscope.

#### 16S rRNA and ITS sequencing and data analysis

16S rRNA and ITS sequencing were performed by Genedenovo Biotechnology. Microbial DNA was extracted from samples using the HiPure Stool DNA Kit (Magen) following the manufacturer’s instructions. Amplicons were assessed by 2% agarose gel electrophoresis and purified using AMPure XP beads (Beckman). Sequencing libraries were prepared using the Illumina DNA Prep Kit (Illumina) according to the manufacturer’s protocol and sequenced on the Novaseq 6000 platform. Raw sequencing reads were quality-filtered and denoised to obtain high-quality representative sequences. Denoising was performed to remove low-quality reads, chimeras, and sequencing errors, generating amplicon sequence variants (ASVs). Representative ASVs were taxonomically classified using a naïve Bayesian classifier via the RDP classifier (version 2.2) against the SILVA database (version 138.1) with a confidence threshold of 0.8. Downstream analyses, including alpha and beta diversity, taxonomic composition, and differential abundance, were conducted using Omicsmart (www.omicsmart.com)

#### RNA sequencing and data analysis

RNA sequencing was performed by Genedenovo Biotechnology. Gastric tissues from *S. cerevisiae*- or *C. albicans*-infected *K19-C2mE* mice were homogenized using bead-beating, and total RNA was extracted with Trizol reagent (Magen) according to the manufacturer’s instructions. RNA quality was assessed prior to library construction. Paired-end (PE) libraries were prepared using the mRNA-seq Lib Prep Kit (RK20308, ABclonal), and library quality was evaluated using an Agilent Bioanalyzer 4150. Sequencing was performed on the MGISEQ-T7 platform with a read length of PE150. Raw sequencing data were processed for quality control and aligned to the reference genome for downstream analyses. Differential gene expression analysis and functional enrichment were conducted using standard bioinformatics pipelines. Immune cell composition in mouse gastric tissues was further inferred using seq-ImmuCC^79^.

#### Untargeted metabolomics and data analysis

Gastric tissue homogenates, serum, and conditioned culture supernatants were collected and stored at −80 °C prior to liquid chromatography–mass spectrometry (LC-MS) analysis at the Core Facility of Biomedical Sciences, Xiamen University. Briefly, 100 μL of each sample was mixed with 400 μL of methanol/acetonitrile (1:1, v/v) and vortexed for 30 s. The mixtures were ultrasonicated for 10 min at 4 °C, incubated at −20 °C for 1 h, and centrifuged at 13,000 rpm for 15 min at 4 °C. Supernatants were transferred to fresh tubes and evaporated to dryness using a vacuum concentrator at 4 °C. Dried samples were reconstituted in 100 μL of acetonitrile/water (1:1, v/v) prior to analysis. Metabolite separation and detection were performed using a Shimadzu Prominence ultra-high-performance liquid chromatography (UHPLC) system (Nexera UHPLC LC-30A) coupled with a TripleTOF 5600^+^ mass spectrometer (SCIEX). Data acquisition was conducted with Analyst TF 1.6 software (SCIEX). Metabolomic data were processed and analyzed using MetaboAnalyst 5.0, employing rigorous deconvolution, normalization, and quality control criteria to ensure high-quality and reliable results.

#### Single-cell RNA-sequencing and data analysis

Single-cell RNA sequencing was performed by Genedenovo Biotechnology (Guangzhou, China). Gastric tissues were collected and dissociated using a Tumor Dissociation Kit (Miltenyi Biotec, Germany) to obtain single-cell suspensions. Cells were encapsulated in gel bead-in-emulsion (GEM) droplets using the GemCode Single-Cell system (10X Genomics) for cDNA library preparation. Libraries were sequenced on an Illumina NovaSeq 6000 platform. Raw sequencing data were processed for quality control, and reads were aligned to the reference genome using Cell Ranger (10x Genomics). Downstream single-cell analyses, including clustering, visualization, and differential expression, were conducted using the “Seurat” R package ^80^ Following initial cell calling, 11,261–19,432 cells were detected per sample. Cells were filtered based on the following quality-control criteria: nFeature_RNA > 200 and < 7,500, nCount_RNA < 100,000, mitochondrial gene percentage (percent.mt) < 20%, and hemoglobin gene percentage (percent.HB) < 5%. After filtering, 8,694–16,675 high-quality cells per sample were retained for normalization, dimensionality reduction, clustering, and cell-type annotation.

#### Flow cytometry

Fresh gastric tissues were rinsed with PBS, minced, and enzymatically digested with trypsin at 37 °C for 10 min. Tissue fragments were further digested in 1 mg/mL collagenase IV (17104019, Thermo Fisher Scientific) for 30 min at 37 °C using a gentle MACS Dissociator to generate a single-cell suspension. Cells were centrifuged at 500 × g for 5 min, resuspended in RPMI 1640, and filtered through a 70 µm MACS SmartStrainer. After a second centrifugation at 500 × g for 5 min, cells were treated with RBC lysis buffer for 5 min at room temperature and resuspended in MACS buffer. Cells were then stained with appropriate antibodies (Live/Dead, CD45, CD11b, F4/80, CD80, CD206, isotype) for flow cytometry. Data were acquired and analyzed using FlowJo software (v. 10.6.2).

#### siRNA Transfection

THP-1, Raw264.7 and BMDMs were seeded in 6-well plates at 50% confluence. Lipofectamine 3000 (L3000015, Thermo Fisher Scientific) was complexed with AR siRNA and incubated for 20 min at room temperature. The siRNA-Lipofectamine mixture was then added dropwise to the cells and incubated for 6 h, after which the medium was replaced with fresh culture medium. Cells were collected for downstream experiments. The siRNA sequences are provided in **Table S4**

#### RT-qPCR

For fungal load analysis, infected gastric tissues were collected and homogenized. For macrophage polarization, THP-1 cells were differentiated with PMA (100 ng/mL) for 24 h and polarized to M1 macrophages with LPS (100 ng/mL) plus IFN-γ (20 ng/mL) for 24 h, or to M2 macrophages with IL-4 (20 ng/mL) for 24 h. Raw264.7 and BMDMs were similarly polarized to M1 or M2 macrophages under the same conditions. Treatments were applied as indicated, including inosine (1mM), ZM241385, or siAR for 24 h. Total RNA from the samples was extracted using the FastPure Cell/Tissue Total RNA Isolation Kit V2 (RC112-01, Vazyme). Reverse transcription was performed using the HiScript III All-in-one RT SuperMix Perfect for qPCR Kit (R333-01, Vazyme), and qPCR was conducted using the Taq Pro Universal SYBR qPCR Master Mix Kit (Q712-02, Vazyme). Primer sequences and detailed information are provided in **Table S4**.

#### Immunofluorescence (IF) Staining

Murine gastric tissue sections (4 µm) were incubated overnight at 4 °C on a shaker with primary antibodies against F4/80 (MCA497G, Bio-Rad), CD80 (ab254579, Abcam), and CD206 (ab64693, Abcam). After washing, sections were incubated with appropriate secondary antibodies-Goat anti-Rat (ab6953, Abcam) or Goat anti-Rabbit (ab150077, Abcam)-for 1 h at room temperature. Nuclei were counterstained and sections were mounted using DAPI-containing anti-fade solution. Fluorescence images were acquired using confocal microscope (Leica) following the manufacturer’s instructions.

#### Inosine and antibody detection

Serum samples from clinical participants were collected and centrifuged at 1,200g for 10 min, followed by a second centrifugation at 12,000g for 10 min. The resulting supernatant was flash-frozen in liquid nitrogen for 1 min and stored at −80□°C until analysis. Inosine levels were measured using the Inosine Quantification Assay Kit (ab126286, Abcam) following the manufacturer’s instructions. Serum IgG against *C. albicans* was quantified using the Anti-human *Candida albicans* IgG ELISA Kit (ab108715, Abcam), and serum antibodies against *H. pylori* were measured using the Anti-human *Helicobacter pylori* Ab ELISA Kit (RD-RX11979, Ruida Henghui Biotech), both according to the respective manufacturer’s protocols.

### QUANTIFICATION AND STATISTICAL ANALYSIS

#### Statistical analysis

Pathological scores and clinical characteristics were compared using the Mann–Whitney U test or Kruskal-Wallis test. Student’s t-test was used to compare means between two groups, while comparisons among multiple groups were performed using one-way analysis of variance (ANOVA). The chi-square (χ²) test was applied to compare the proportions of mice with dysplasia or lymph node metastasis between groups. Survival analyses were conducted using the log-rank (Mantel–Cox) test. All statistical analyses were performed using GraphPad Prism 8.0.1. A P value < 0.05 was considered statistically significant. Detailed information regarding statistical tests and sample sizes is provided in the corresponding figure legends.

**Figure S1.**
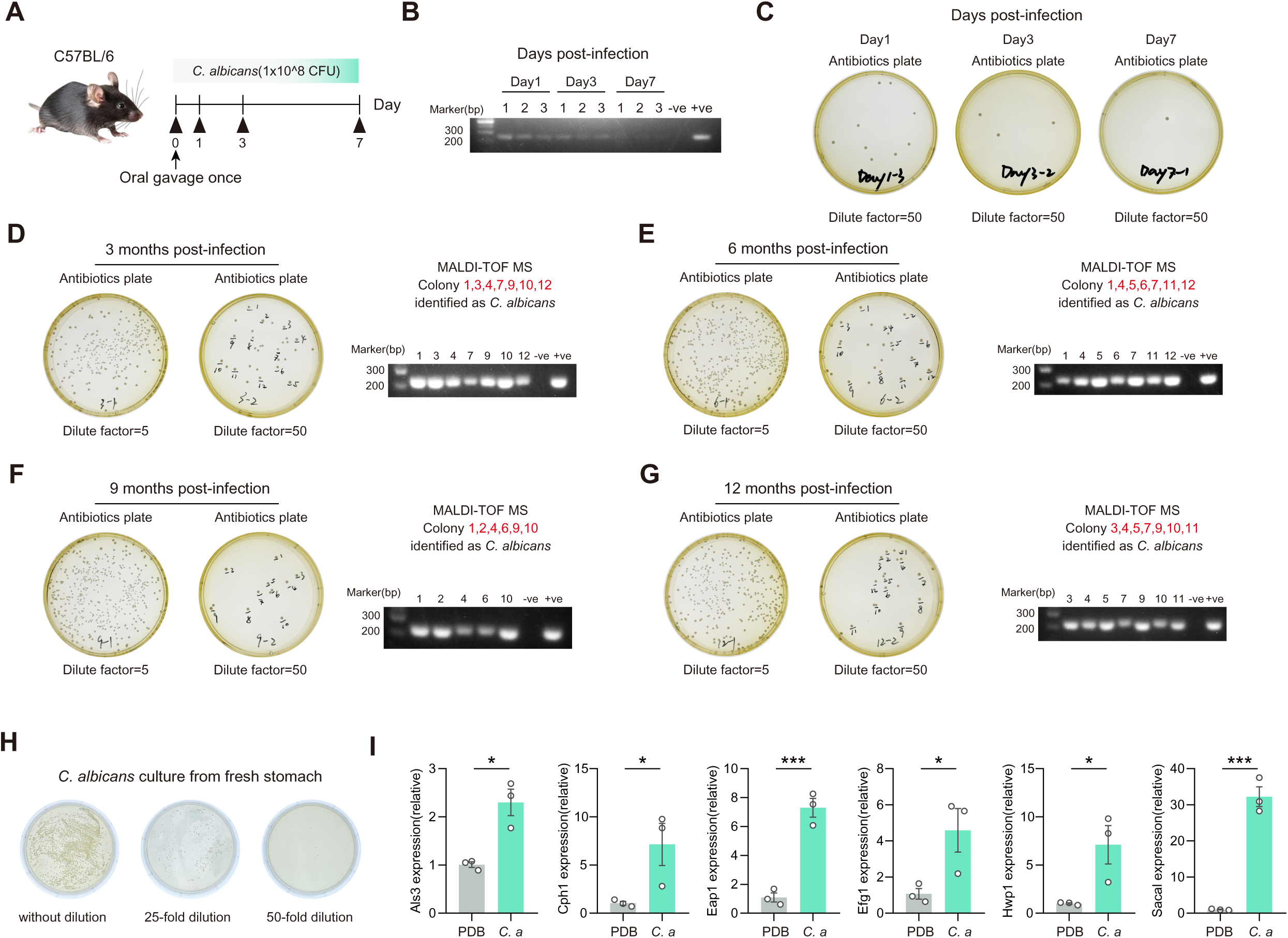
*Candida albicans* colonization of the gastric mucosa following long-term infection in mice. (A and B) PCR analysis of *C. albicans* colonization in gastric tissue of C57BL/6 male mice at 1, 3, and 7 days after a single oral gavage (n =3/time point) (C) Stomach fungal culture of *C. albicans*-infected mice at 1, 3, and 7 days after a single oral gavage. (D-G) Stomach fungal culture at 3, 6, 9, and 12 months post-C. albicans infection (panels D-G, respectively), with colony PCR verification of MALDI-TOF MS identifications (Table 1) (H-I) Stomach fungal culture of *C. albicans* (H) and the relative mRNA expression of *C. albicans* (I) in germ-free mice infected with *C. albicans*. Data were presented as mean ± SEM. Each point represents one subject. Student’s t-test (I) was used to assess the statistical significance between groups, * p < 0.05, *** p < 0.001.

**Figure S2.**
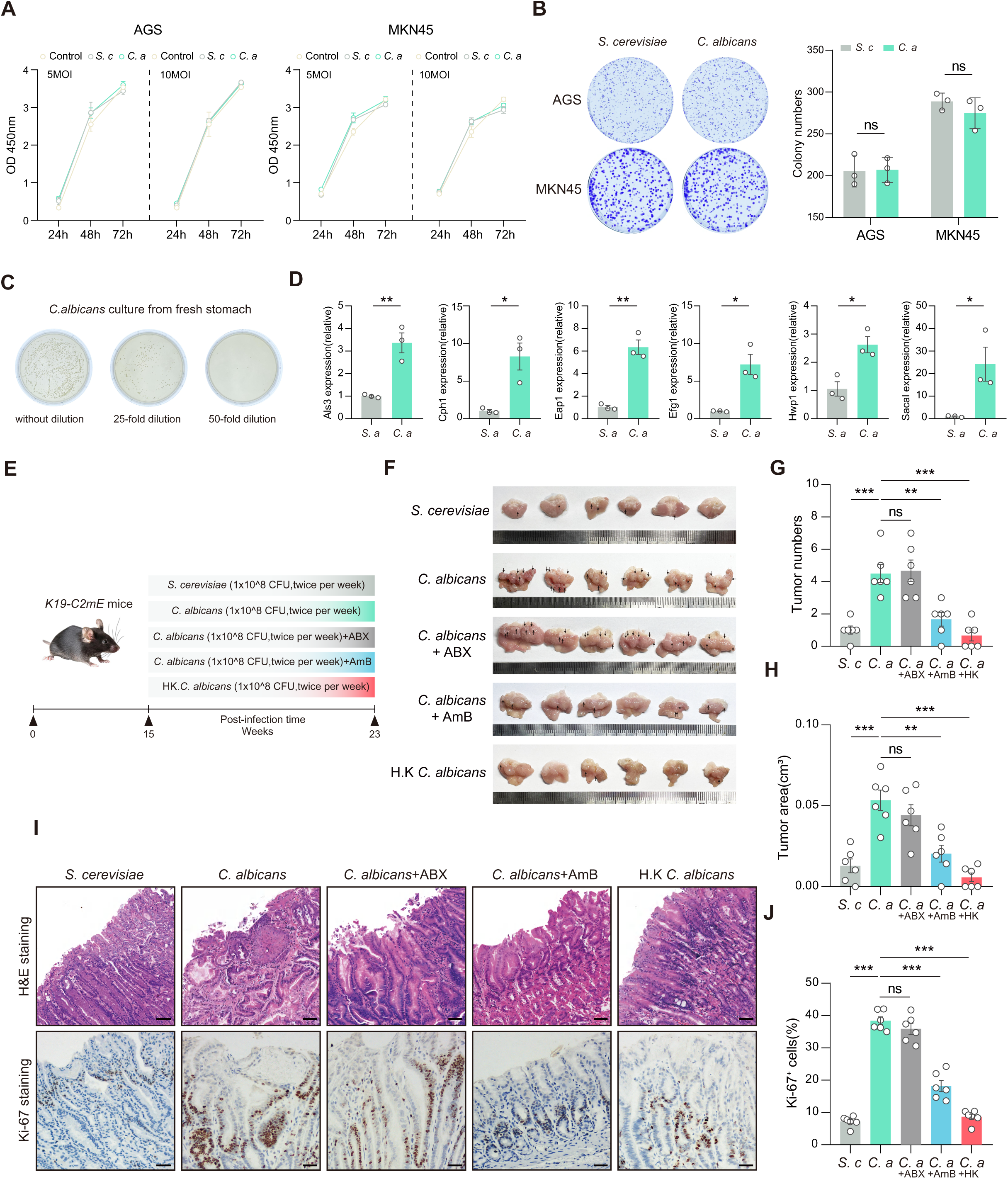
The presence of viable *C. albicans* within the tumor is essential and solely sufficient to promote protumor effects. (A-B) CCK-8(A) and Colony(B) formation assay of gastric cancer cells co-cultured with *S.cerevisiae* or *C.albicans* at 5MOI or 10MOI. (C-D) Stomach fungal culture of *C. albicans* (C) and the relative mRNA expression of *C. albicans* (D) in *K19-C2mE* mice infected with *C. albicans*. (E) *K19-C2mE* mice were orally administered *S. cerevisiae*, *C. albicans*, IG with AmB or ABX, and HK *C. albicans* for 2 consecutive months (n = 6/group). HK *C. albicans*, heat killing C. albicans under 95℃ for 2h; IG, intragastric administration; AmB: Amphotericin B; ABX, vancomycin, neomycin, ampicillin, and metronidazole. (F) Representative morphology images of stomachs in the indicated groups; scale bars, 5 mm. (G-H) Quantification of tumor numbers(G) and tumor area (H) in the indicated groups. (I) Representative H&E-stained and Ki-67-stained images of stomach sections from indicated groups; scale bars, 100μm. (J) Quantification of Ki-67 proliferation rates in the indicated groups. Data were presented as mean ± SEM. Each point represents one subject. Student’s t-test (B and D), Kruskal-Wallis test (G) and One-way ANOVA (H and J) were used to assess the statistical significance between groups, ns, nonsignificant, * p < 0.05, ** p < 0.01, *** p < 0.001.

**Figure S3.**
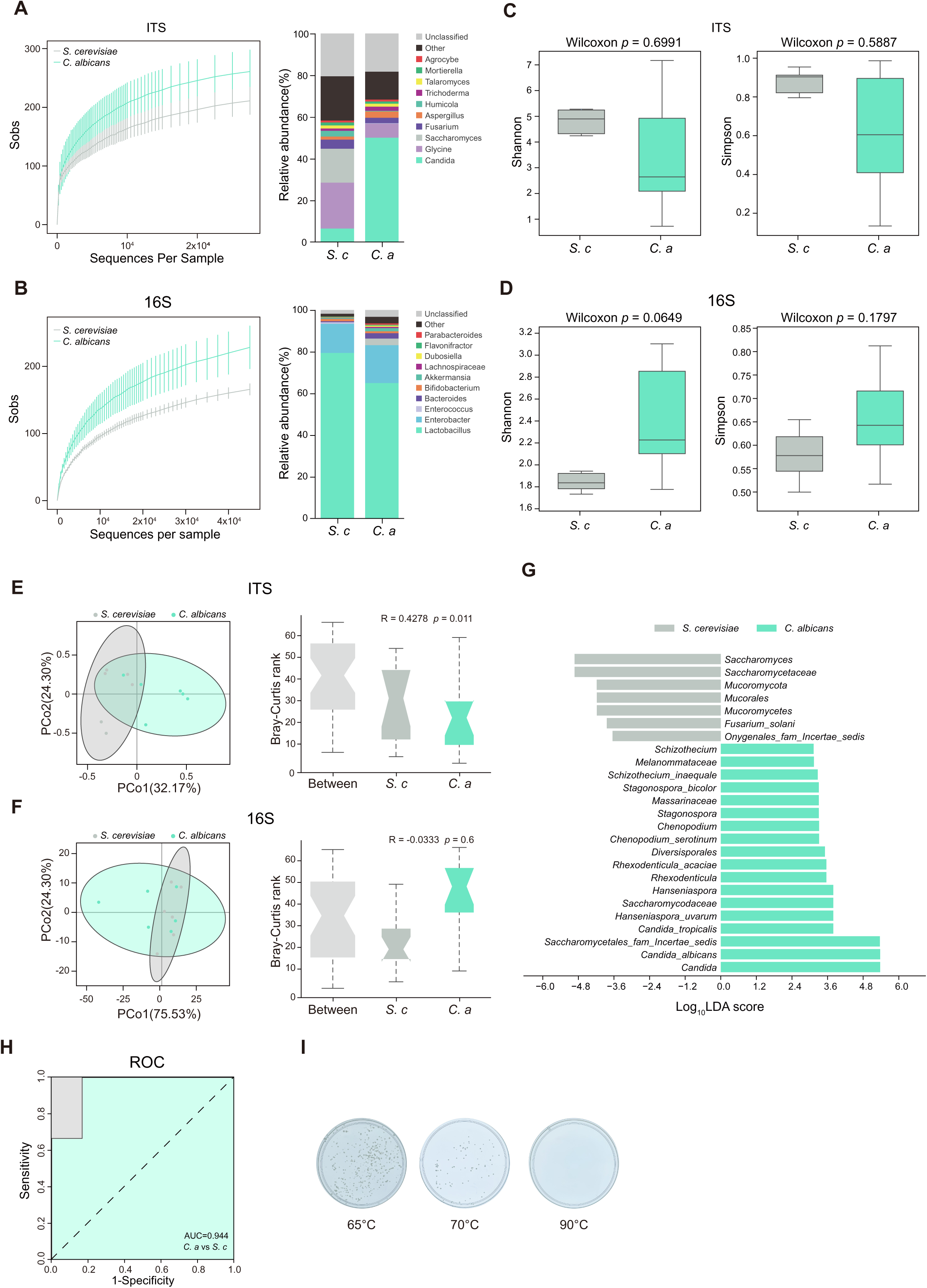
The presence of viable *C. albicans* within the tumor is crucial and independently sufficient to induce protumor effects. (A) Species accumulation curve (left) and Fungi at Genu taxonomic levels (right) of *S.cerevisiae* or *C.albicans* infected *K19-C2mE* mice by ITS sequencing. (B) Species accumulation curve (left) and Fungi at Genu taxonomic levels (right) of *S.cerevisiae* or *C.albicans* infected *K19-C2mE* mice by 16S sequencing. (C) Alpha diversity of *S.cerevisiae* or *C. albicans* infected *K19-C2mE* mice by ITS sequencing. (D) Alpha diversity of *S.cerevisiae* or *C. albicans* infected *K19-C2mE* mice by 16S sequencing. (E) PCoA based on Bray-Curtis distances to assess beta diversity of ITS sequencing. (F) PCoA based on Bray-Curtis distances to assess beta diversity of 16S sequencing. (G) LDA scores identify features with varying abundances between *S. cerevisiae* and *C. albicans*-infected groups. Features with an LDA score >2.5 are considered significantly differentiating between the groups. (H) ROC analysis using genus scores to predict *C. albicans*. ROC, receiver operating characteristics (I) Growth of *Candida albicans* at various heat-killing temperature. Data were presented as mean ± SEM.Each point represents one subject. Wilcoxon t test (C, D, E, and F) was used to assess the statistical significance between groups.

**Figure S4.**
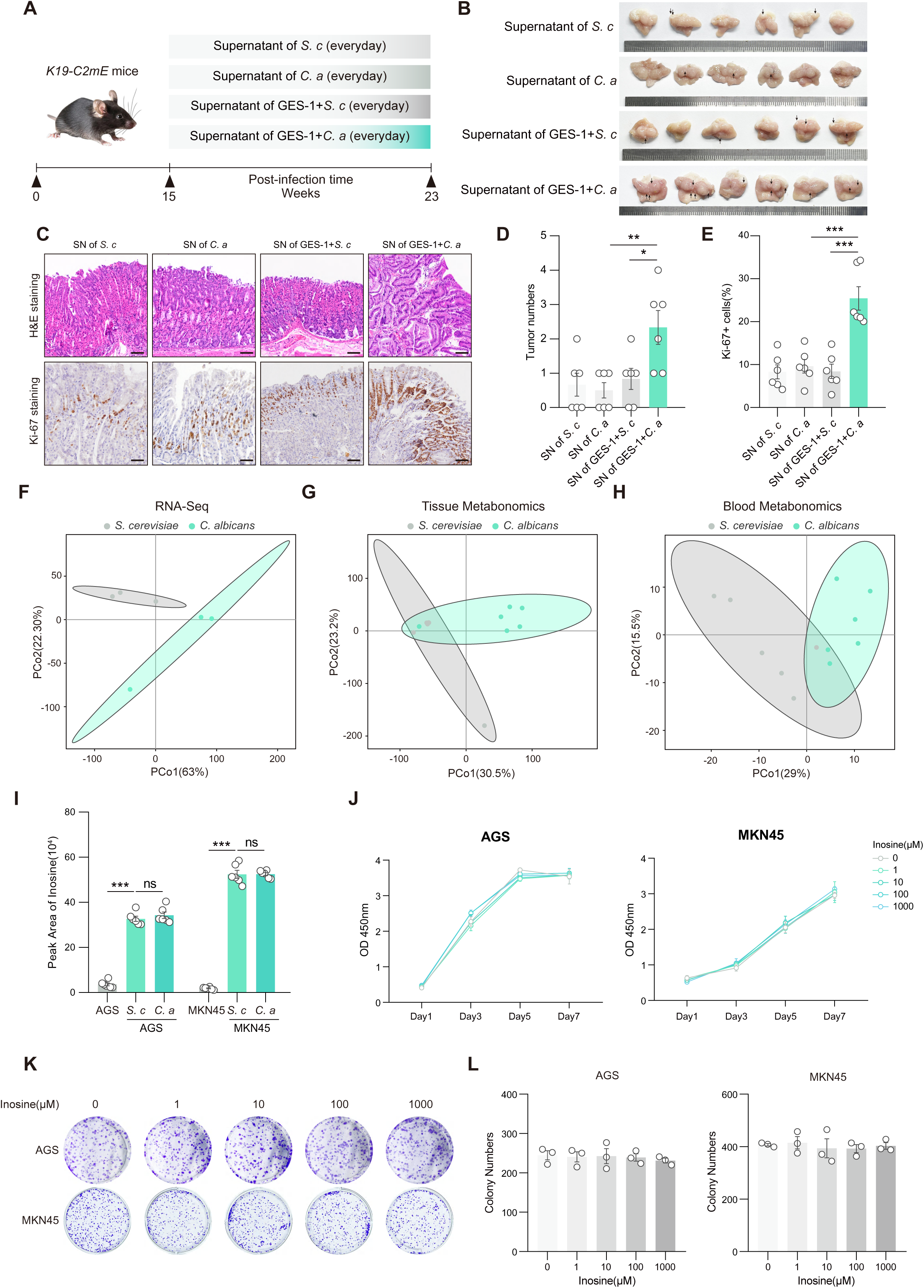
*Candida albicans* enhances inosine secretion to promote gastric tumorigenesis. (A) *K19-C2mE* mice were orally administered supernatant of *S. cerevisiae* supernatant of *C. albicans*, Supernatant of *S. cerevisiae* co-coluture with GES-1, Supernatant of *C. albicans* co-coluture with GES-1 for 2 consecutive months (n = 6/group) (B) Representative morphology images of stomachs in the indicated groups; scale bars, 5 mm. (C) Representative H&E-stained and Ki-67-stained images of stomach sections from indicated groups; scale bars, 100μm. (D-E) Quantification of tumor numbers (D) and Ki-67 proliferation rates(E) in the indicated groups. (F) PCoA of transcriptomics on gastric tissues from *S.cerevisiae* or *C. albicans* infected *K19-C2mE* mice (n = 3/group). (G) PCoA of metabolomics on gastric tissues from *S.cerevisiae* or *C. albicans* infected *K19-C2mE* mice (n = 6/group). (H) PCoA of metabolomics on serum from *S.cerevisiae* or *C. albicans* infected *K19-C2mE* mice (n = 6/group). (I) The relative concentration of inosine in different treatment groups in AGS and MKN45 cells. (J-L) CCK-8 (J) and colony formation (K and L) assay of AGS and MKN45 cells treated with different concentrations of inosine. Data were presented as mean ± SEM. Each point represents one subject. Kruskal-Wallis test (D) and One-way ANOVA (E, I, J and L) were used to assess the statistical significance between groups, ns, nonsignificant, * p < 0.05, ** p < 0.01, *** p < 0.001.

**Figure S5.**
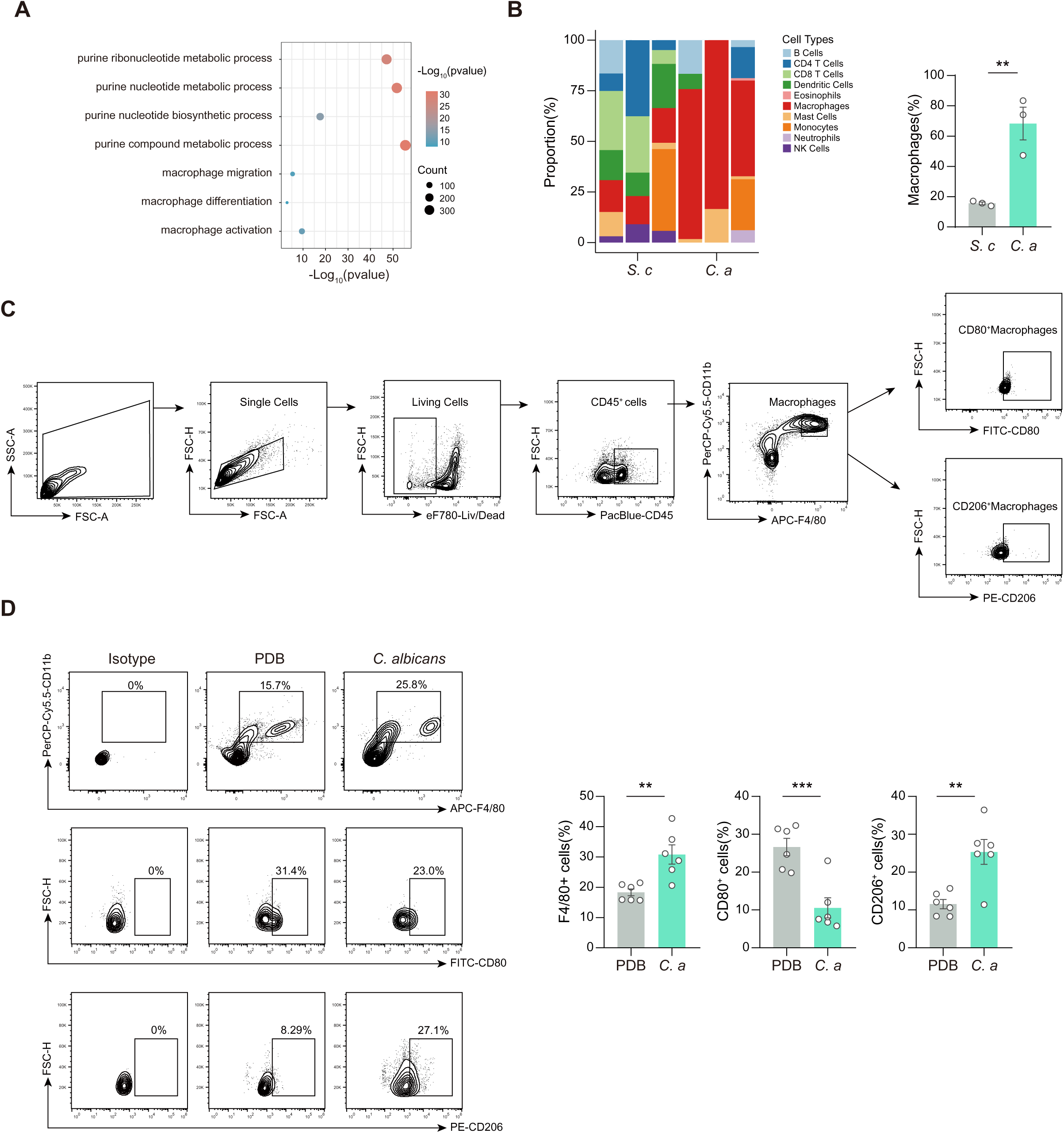
*C.albicans* modifies M2 macrophage activity to drive tumor growth. (A) Differentially expressed genes were analyzed using GO enrichment for biological processes. (B-C) Seq-immuCC analysis of RNA sequencing to predict the immune cell composition in *C. albicans* infected *K19-C2mE* mice. (D) Flow cytometry gating strategy for macrophage analysis (E) Flow cytometry analysis of M1 or M2 macrophage (CD80 or CD163) proportions between PDB and *C. albicans*-infected germ-free mice. Data were presented as mean ± SEM. Each point represents one subject. Student’s t-test (B, and D) was used to assess the statistical significance between groups, ** p < 0.01, *** p < 0.001.

**Figure S6.**
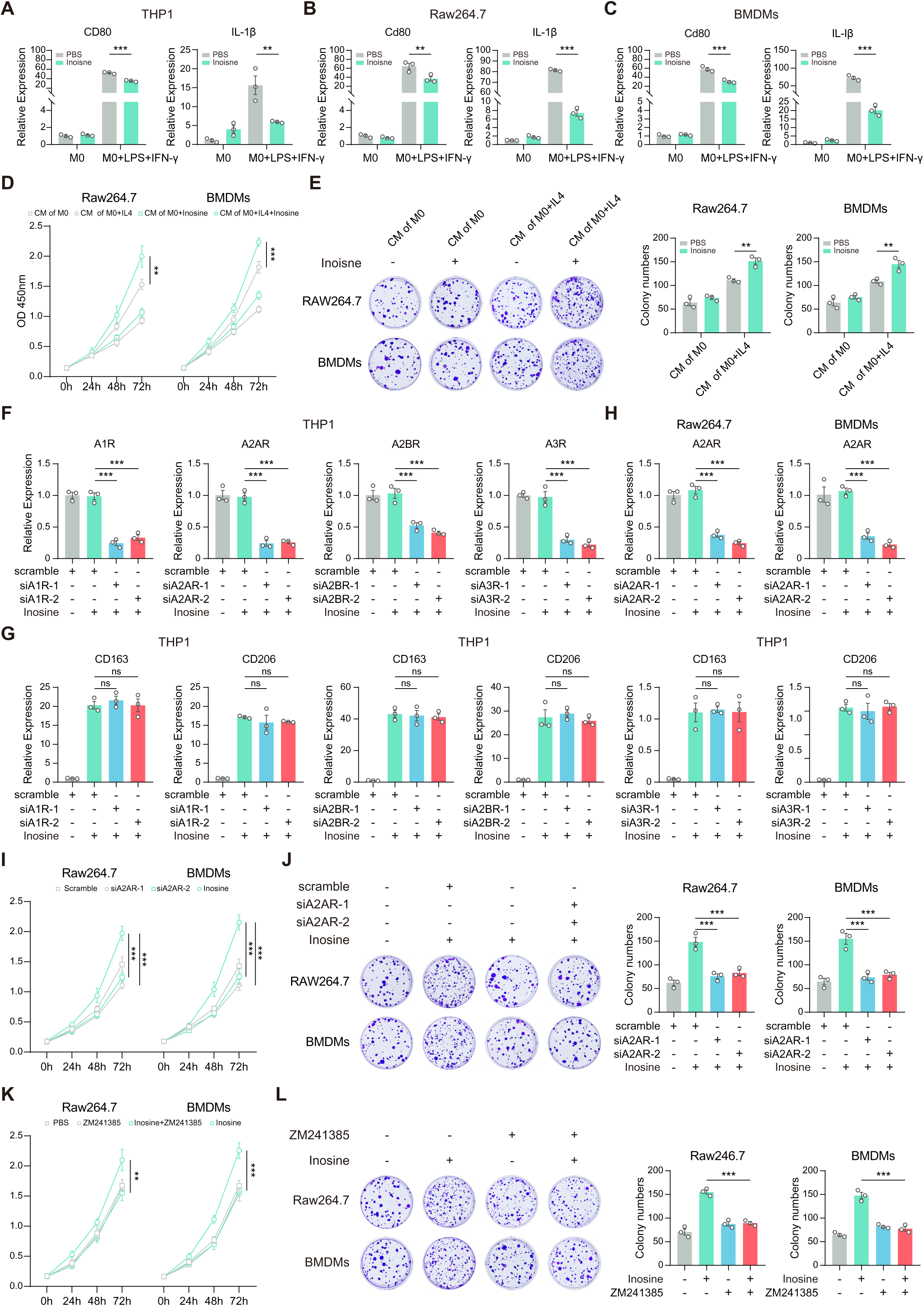
Inosine activates A2AR receptors in macrophages to trigger tumorigenesis. (A-C) RT-qPCR analysis of M1 markers (CD80 and IL-1β) in THP-1 (A), RAW264.7 (B), and bone marrow-derived macrophages (BMDMs) (C) treated with or without inosine (1 mM) for 24 hours under M2-polarizing conditions. (D-E) CCK-8 assay (D) and colony formation assay (E) assessing the effects of conditioned media from differentially polarized-RAW264.7 and BMDMs on ATK cell proliferation. (F) RT-qPCR analysis of adenosine receptor (AR) family expression in THP-1 cells transfected with scramble, A1R-targeting siRNA (siA1R), A2AR-targeting siRNA (siA2AR), A2BR-targeting siRNA (siA2BR), or A3R-targeting siRNA (siA3R). (G) RT-qPCR analysis of CD206 and CD163 expression to assess the role of AR family in inosine-mediated effects. THP-1 cells were transfected with scramble, A1R-targeting siRNA (siA1R), A2AR-targeting siRNA (siA2AR), A2BR-targeting siRNA (siA2BR), or A3R-targeting siRNA (siA3R) and treated with or without inosine (1 mM) for 24 hours under M2-polarizing conditions. (H) RT-qPCR analysis of A2AR expression in RAW264.7 and BMDMs transfected with scramble or A2AR-targeting siRNA (siA2AR). (I-J) CCK-8 assay (I) and colony formation assay (J) assessing the effects of conditioned media from differentially polarized macrophages, with or without A2AR silencing, on tumor cell proliferation. (K-L) CCK-8 assay (K) and colony formation assay (L) assessing the effects of conditioned media from differentially polarized macrophages, with or without ZM241385 (5µM), on tumor cell proliferation. Data were presented as mean ± SEM. Each point represents one subject. One-way ANOVA (A, B, C, D, E, F, G, H, I, J, K, and L) was used to assess the statistical significance between groups, ns, nonsignificant, ** p < 0.01, *** p < 0.001.

**Figure S7.**
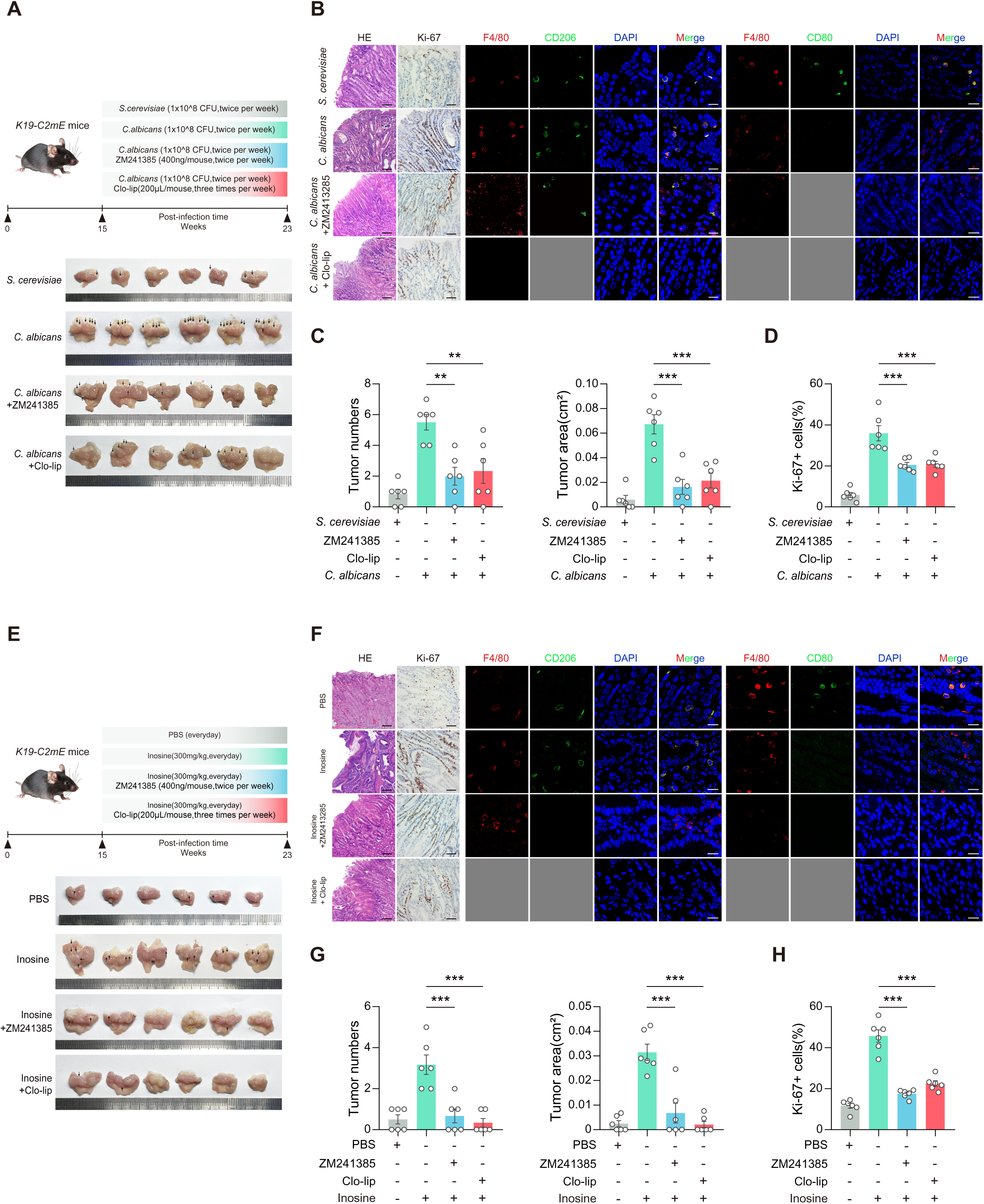
*C. albicans* modulates macrophage pro-carcinogenic effects via the inosine-A2AR axis. (A) Representative stomach images from *K19-C2mE* mice after 2-month oral administration of *S. cerevisiae* or *C. albicans*, with or without co-treatment with ZM241385 (0.4 μg per mouse, IP) or clodronate liposomes (200 μL per mouse, IV) (n=6/group). (B) Representative H&E-stained and Ki-67-stained images of stomach sections from indicated groups; scale bars, 100μm. (C-D) Quantification of tumor numbers, tumor area (C) and Ki-67 proliferation rates(D) in the indicated groups. (E) Representative stomach images from *K19-C2mE* mice after 2-month oral administration of PBS or inosine, with or without co-treatment with ZM241385 (0.4 μg per mouse, IP) or clodronate liposomes (200 μL per mouse, IV) (n=6/group). (F) Representative H&E-stained and Ki-67-stained images of stomach sections from indicated groups; scale bars, 100μm. (G-H) Quantification of tumor numbers, tumor area (G) and Ki-67 proliferation rates(H) in the indicated groups. Data were presented as mean ± SEM. Each point represents one subject. Kruskal-Wallis test (C/tumor numbers and G/tumor numbers) and One-way ANOVA (C/tumor area, D, G/tumor area, and H) were used to assess the statistical significance between groups, ** p < 0.01, *** p < 0.001.

